# Ups and downs of liquid-liquid transitions in GUV membranes from osmolarity and aspiration tensions

**DOI:** 10.64898/2026.09.03.748981

**Authors:** Takashi Kuroyanagi, Kent J. Wilson, Aymeric Chorlay, Caitlin E. Cornell, Daniel A. Fletcher, Sarah L. Keller

## Abstract

Lipid membranes undergo liquid-liquid phase separation to form micron-scale domains. Several physical parameters affect this phase transition. For example, a decrease in temperature causes membranes to demix, as does an increase in hydrostatic pressure. However, measurements to determine how tension affects membrane phase separation have yielded conflicting results. Experiments that have applied osmotic pressure differences to a population of vesicles have reported an increase in the membrane’s miscibility transition temperature. Conversely, experiments that have applied micropipette aspiration or substrate stretching to single membranes have reported a decrease. Here, we find that both osmotic pressure and micropipette aspiration can increase transition temperatures. We discuss how membrane pores and hidden areas in membranes present challenges to researchers seeking to quantitatively convert experimentally measured osmolarity differences into membrane tensions. We show that challenges of comparing data from different osmotic pressure experiments can be mitigated by renormalizing osmolarity differences, specifically by dividing by the exterior osmolarity. We discuss our results in the context of existing theoretical predictions and in light of four known effects of increasing tension on vesicle membranes: 1) a reduction in hidden area of tubes and aggregates, 2) a reduction in out-of-plane thermal fluctuations, 3) an increase in the area per lipid, and 4) the formation of pores. First, hidden area in tubes and other structures explains why vesicles appear to sustain high osmolarity differences. Next, changes in a membrane’s fluctuations and molecular area may explain shifts in miscibility transition temperatures. Finally, pores can explain time dependences. Overall, our results highlight the need for new theory, simulation, and experiments that unify predictions of how transition temperatures vary over all four regimes of membrane tension.

**STATEMENT OF SIGNIFICANCE:** Liquid-liquid phase separation occurs in artificial lipid membranes and in some natural membranes. Although there is consensus that applying tension to membranes should affect this phase transition, it is unclear whether tension should induce or suppress membrane phase separation, even in artificial membranes. Here, we show that two experimental methods of applying tension cause liquid domains to appear in membranes at constant temperature. We consider four different regimes of membrane tension. We introduce two speculative scenarios in which tension has nonmonotonic effects on membrane phase separation, and we evaluate the scenarios in context of current theory and experiments. We find that a scenario in which tension enhances and then suppresses phase separation is consistent with most (but not all) results.

## INTRODUCTION

Cells are known to exhibit spatial and temporal variations in membrane tension during fibroblast spreading (1), cell migration (2, 3), and metastasis (4). Likewise, cells have evolved a variety of ways to adjust membrane tension. For example, in yeast, a key function of the vacuole (an organelle) is to prevent lysis when high tension arises due to an imbalance of osmotic pressures across the plasma membrane (5). Membranes of yeast vacuoles are also notable because they undergo liquid-liquid phase separation (6), a transition that is sensitive to environmental conditions (7, 8).

**Figure 1:**
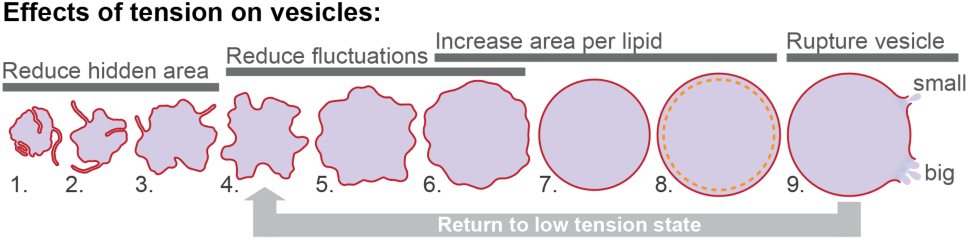
Four effects of membrane tension on lipid vesicles. At very low tensions, tubes and lipid aggregates are pulled into the membrane. As tension increases, thermal fluctuations diminish, and the lateral area per lipid increases. Throughout, pores may form. Small pores can reseal, whereas larger ruptures can expel fluid and relieve membrane tension.

Tension is known to shift membrane miscibility temperatures (*T*_mix_), at least in artificial membranes (9–15). One exciting implication is that cells could use spatial and temporal variations in tension to drive transient phase transitions. However, major challenges persist in predicting whether tension increases or decreases *T*_mix_, in testing predictions in the context of experimental measurements, or even in comparing between experiments. Some of these challenges arise because tension has four known effects on vesicle membranes (Figure 1).

### Reduction of hidden area

At the lowest tensions, “hidden area” in tubes or lipid aggregates is retracted into the membrane (16, 17). Tensions in this regime are so low that they are often not measurable. The amount of hidden area varies widely, from 2% of total membrane area (in vesicles made by gentle hydration from PC lipids (17)) to 20-50% of the area (in vesicles made by electroformation from POPC lipids with 2-4 mol% GM1 lipids in the inner leaflet (18)). Tubes with high curvature can produce spontaneous tensions of ∼10^−2^ mN/m, whereas the vesicle’s mechanical tension is on the order of 10^−4^ mN/m (18).

### Reduction of fluctuations

Out-of-plane thermal fluctuations of the membrane exist at all non-zero tensions (19). For tensions from 10^−5^ to ∼0.5 mN/m, increases in the observed vesicle area are dominated by decreases in thermal fluctuations (19, 20).

### Stretching

Beyond tensions of ∼0.5 to 1 mN/m, increases in observed vesicle area are dominated by stretching of the membrane to increase the lateral area per lipid (19, 21). This effect persists to some extent over all non-zero tensions (19).

### Formation of pores and membrane rupture

Throughout, transient pores stochastically open and close, which allow the vesicle’s internal solution to leak out, resetting the tension at a lower value (e.g., (13, 22) and references therein). Pore size is proportional to the solution viscosity; pores are submicron in low viscosity solutions like water and sucrose (22). Membrane rupture is irreversible at tensions that exceed the lysis tension, which varies with lipid composition: for membranes of pure unsaturated or mono-unsaturated lipids, the lysis tension is ∼10 mN/m, whereas for polyunsaturated lipids and eggPC lipids, it ranges from 3-5 mN/m (21, 23), and references therein. These lysis tensions correspond to vesicles that have experienced an increase in surface area of ≤ 4% from when vesicles first appeared spherical (21, 23).

Two common methods of applying tension to membranes are micropipette aspiration and the application of osmotic pressure. Strengths of the micropipette approach are that the experiments are conceptually straightforward, and that well-established theory enables researchers to directly extract membrane tension from images (21). A disadvantage is that micropipette experiments are challenging, require specialized instrumentation, and are low throughput because they measure only one vesicle at a time (11). In contrast, applying osmotic pressure across membranes is easy to implement over a large population of vesicles, but variances are large, and results can be difficult to interpret (9, 10, 12, 13, 24, 25).

Micropipette aspiration and osmotic pressure have both been used to assess how tension affects liquid-liquid phase separation of membranes. However, until now, the two techniques have been used to different ends. To our knowledge, micropipette aspiration has previously been used to measure shifts in membrane miscibility temperatures (*T*_mix_) only in the stretching regime at high tensions, where thermal fluctuations are smaller (11). In contrast, osmotic pressure has been applied over all tension regimes, including throughout cycles of vesicle rupture, which cause liquid domains to appear and disappear in membranes (9, 10, 12, 24, 25) (see also Supplemental Movies S1 and S2).

**TABLE 1.** Tension makes domains appear.

| Experimental observation | How tension applied | System | Refs |
|---|---|---|---|
| **Tension makes domains appear** |  |  |  |
| Increase *T*<sub>mix</sub> of Lo/Ld phases | Osmotic pressure | 3-component GUVs* | (9, 10, 12, 24, 25) |
| Increase *T*<sub>mix</sub> of S/L phases | Osmotic pressure | 2-component GUVs† | (10, 24) |
| **Tension makes domains disappear** |  |  |  |
| Experimental observation | How tension applied | System | Refs |
| Decrease *T*<sub>mix</sub> of Lo/Ld phases | Micropipette aspiration | 3-component GUVs* | (11) |
| Decrease *T*<sub>mix</sub> of Lo/Ld phases | Substrate stretching | 3-component SLBs* | (26) |
| Decrease *T*<sub>mix</sub> of Lo/Ld phases | Hydrostatic pressure | 3-component GUVs* | (27–30) |
| Decrease *T*<sub>mix</sub> of S/L phases | Micropipette aspiration | 2-component GUVs† | (31) |

\* GUVs (giant unilamellar vesicles) or SLBs (supported lipid bilayers) were made from ternary mixtures of a lipid with a high melting temperature (DMPC, DPPC, eggSM, or brainSM), a lipid with a low melting temperature (DOPC, POPC, or DiPhyPC) and cholesterol.
† GUVs were made from binary mixtures of a lipid with a high melting temperature (DPPC) and a lipid with a low melting temperature (DOPC).

The scientific literature contains well-known results regarding how osmotic pressure affects vesicles. By holding vesicles a few degrees above *T*_mix_ (such that their membranes are uniformly mixed) and swelling them, it is straightforward to observe that liquid domains appear, which indicates an *increase* in *T*_mix_ (9, 10, 12, 24, 25). Osmotic pressure differences are typically imposed by forming vesicles in a sugar solution and then diluting the exterior solution. If the difference in osmolarities is big enough, a large pore forms, which relieves tension, decreases *T*_mix_, and causes an abrupt decrease in vesicle radius, sometimes cyclically (9, 10, 12, 13, 22, 24, 25).

Our interest in membrane tension is motivated by several perplexing inconsistencies in the literature (Table 1) that we attempt to resolve in this manuscript, not always conclusively. How can membranes in osmotic pressure experiments appear to withstand very high tensions (11), whereas micropipette experiments measure lower lysis tensions (22, 24)? How can *increases* in *T*_mix_ observed in osmotic pressure experiments (10, 11, 13, 25, 26) be consistent with *decreases* in *T*_mix_ predicted by theory based on the Gibbs-Duhem equation (16) and observed in experiments using micropipette aspiration of GUVs, stretching of supported bilayers, and hydrostatic pressure (12, 27–31)? Given that the apparent inconsistencies persist in membranes with solid phases, are the differences due to only the choice of experimental technique (10, 31)?

Here, we measure shifts in liquid-liquid phase transitions in vesicle membranes under tension. We apply tension both through differences in solution osmolarity (Δ*c*) and through micropipette aspiration at constant temperature. Based on our results, we introduce two speculative scenarios in which miscibility temperatures of membranes could vary nonmonotonically with tension.

We also quantify the magnitude of systematic errors that we (and others) introduce when we apply osmotic pressure differences across vesicles. Small osmotic pressure differences are commonly applied to reduce excess area and enable phase-separated domains to freely diffuse and coalesce in membranes, as in (33, 34). These osmotic differences have the potential to shift membrane miscibility temperatures.

## MATERIALS AND METHODS

### Chemicals

The lipids dipalmitoyl(16:0)-phosphocholine (DPPC), diphytanoyl(16:0-Me)-phosphocholine (DiPhyPC), dimyristoyl(14:0)-glycerophosphocholine (DMPC), ditridecanoyl(13:0)-glycerophosphocholine, dipalmitoyl(16:0)-glycerophosphoserine (DPPS), dipalmitoyl(16:0)-phosphoethanolamine-N-(lissamine rhodamine B sulfonyl) (Rhod-DPPE) and dioleoyl(18:1)-glycerophosphoethanolamine-N-(lissamine rhodamine B sulfonyl) (Rhod-DOPE) were purchased from Avanti Polar Lipids (Alabaster, AL) and were used without further purification (quoted at 99%) at their supplied nominal concentrations. Cholesterol was purchased from Sigma-Aldrich in powder form and was dissolved in chloroform at 10 mg/mL or 2 mg/mL. Osmotic solutions were made by dissolving either sucrose (ACS grade from Fisher Chemical) or trimethylamine-N-oxide (TMAO, 95% purity from Sigma-Aldrich) in 18 MΩ-cm water; structures are in Supplemental Figure S1). In leakage experiments, calcein (powder form, Sigma-Aldrich) was added to samples at ∼50 µM in water. Vacuum grease (MOLYKOTE high vacuum grease) was from DuPont. All chemicals were used as supplied, without further purification.

### Electroformation

Each sample began as a mixture of 0.25 mg of lipids in chloroform. Composition 1 was a 40/20/40 mixture of DiPhyPC/DPPC/chol, composition 2 was a 20/40/40 mixture, composition 3 was a 33/17/50 mixture, and composition 4 was a 42.5/15/42.5 mixture. In samples of compositions 1 and 2, 0.8 mol% Rhod-DPPE was added to fluorescently label the liquid-disordered phase, and in samples of compositions 3 and 4, 0.8 mol% Rhod-DOPE (the unsaturated version) was used. A glass pipette was used to deposit the sample on a glass slide coated with indium-tin-oxide (Delta Technologies, Loveland, CO) at 60°C. Then the long side of the pipette was passed back and forth across the surface to spread a thin, even film of lipid across an area of ∼1.5 x 3.0 cm as the chloroform dried. To evaporate residual chloroform from the film, the slide was placed in a vacuum desiccator for ≥30 min. Two 1-mm-thick Teflon spacers were affixed near the edges of the slide with vacuum grease, and a second ITO-coated slide assembled face-down on the spacers to make a capacitive chamber. The chamber was filled with osmotic solution, sealed with additional vacuum grease, and placed in an oven 60°C. This temperature is well above the highest melting temperature of all phospholipids (32). An AC voltage of 1 V at 10 Hz was applied across the ITO surfaces for 1 hour at 60 °C to create a solution of giant unilamellar vesicles. For micropipette experiments, vesicles were made in a nearly identical manner, using a 0.3-mm rubber spacer and an AC voltage of 1 V at 10 Hz, at a temperature of 55 °C (33).

### Osmotic pressure

Osmotic pressures calculated from π = *RTc* (where *R* is the gas constant, *T* is temperature, and *c* is concentration) were validated to be within 5% of directly measured values (Wescor VAPRO 5520 vapor pressure osmometer) for a series of sucrose solutions. After electroformation, the osmotic pressure inside and outside every vesicle was putatively equal.

The vesicle solution was poured from the electroformation chamber into an Eppendorf tube. Dilution was performed in two steps. In the first step, the vesicle solution was diluted with three volumes of the same isotonic solution used for electroformation, and the Eppendorf tube was gently inverted three times to mix the solutions. Directly before imaging, the second dilution step was performed. In the second step, aliquots (either 30 µL or 60 µL) of the diluted vesicle solution were transferred to new Eppendorf tubes, and each was further diluted with of a second solution (either 270 µL or 240 µL, respectively, for a total volume of 300 μL). For control samples, the second solution was the same isotonic solution used for electroformation. For test samples, the second solution had a lower concentration of osmolyte. The quoted concentration of the osmolyte outside the vesicles (*c*_outside_) accounts for both the 60 µL of initial solution and the 240 µL of diluting solution, but it neglects the volume inside vesicles. To mix solutions after the second dilution step, the tip of a plastic pipette was cut to make a hole of 3 mm diameter, and the solution was gently pipetted three times. Fewer pipette cycles resulted in higher sample-to-sample variability. More pipette cycles sometimes resulted in smaller shifts in *T*_mix_ values, likely because vesicles were rupturing under shear.

### Micropipette aspiration

Aspiration was carried out as previously described (33). In brief, a filament puller (Sutter Instruments) drew pipettes from borosilicate glass capillaries (G100-4 from Harvard Apparatus: 1.0 mm outer diameter, 0.58 mm inner diameter, 100 mm length). A microforge (MicroData Instruments) smoothed the ∼5 µm diameter pipette opening. Pipettes were filled for ≥30 min with 100 mM sucrose, then for 15 min with 10% bovine serum albumin solution to passivate the micropipette surface. Pipettes were mounted on a TransferMan4r manipulator (Eppendorf) on a confocal microscope (Nikon Eclipse Ti2) and attached to a syringe pump (CellTram, Eppendorf). Vesicles were placed in a chamber on the microscope stage filled with ≥1 mL of solution. This solution was in contact with air for ≤10 min, after which a new vesicle solution was used. The chamber temperature was set by an Okolab Uno stagetop incubator, and sample temperature was measured using a FLIR C5 infrared camera. Membrane tension was found by plugging the suction pressure (Δ*P*_suction_, measured by sensor DP103, Validyne Engineering) and the radii of the pipette (*R*_pipette_) and vesicle (*R*_vesicle_) into the Laplace equation: tension = Δ*P*_suction_/[2(*R*_pipette_^-1^ – *R*_vesicle_^-1^)].

### Imaging

Within 5 min of the second dilution step (and within 48 h. of electroformation), each vesicle solution was sandwiched between coverslips with a seal of vacuum grease, placed on a home-built temperature stage, and imaged with a Hamamatsu C13440 digital camera on a Nikon Eclipse ME600L upright epifluorescence microscope.

### Measuring and calculating error in *T*_mix_

Vesicle populations were heated from ∼20 °C to 50 °C in steps of 1 °C near the miscibility transition temperature (*T*_mix_), and steps of 5 °C otherwise. A single user manually recorded the percent of phase-separated vesicles for at least 100 vesicles at each temperature. This process typically took ∼1 hour. Only vesicles with diameters greater than 20 μm were counted because reliably distinguishing phase separation in smaller vesicles was difficult. *T*_mix_ was found from a fit of a nonlinear least squares regression to the following sigmoidal curve: *Percent phase-separated* = *Max* [1 – (1 + exp (–(*T* – *T*_mix_)/*w*))^−1^], where *Max* is the maximum percent of phase-separated vesicles, *T* is temperature, and *w* is the width of the sigmoidal curve. For each fit of the sigmoidal curve, the standard error was obtained from the diagonal element of the parameter covariance matrix. The half-width, *h*, of the approximate two-sided 95% confidence interval was calculated using the *t-*distribution with *n* – 3 degrees of freedom, where *n* is the number of data points, accounting for the three parameters used to fit the sigmoidal curve. Vertical error bars in *T*_mix_ for vesicles in isoomotic solutions (Δ*c* = *c*_inside_ – *c*_outside_ = 0) are estimated as ±*h.* Vertical error bars in Δ*T*_mix_ for Δ*c* ≠ 0 are estimated assuming independent fits as: [(*h*_Δ*c≠*0_)^2^ + (*h*_Δ*c*=0_)^2^]^½^. Horizontal error bars in Δ*T*_mix_ for Δ*c* ≠ 0 represent a 5% relative uncertainty in sucrose concentrations, the maximum discrepancy between calculated osmolarities and measured osmolarities.

## RESULTS

Liquid-liquid phase separation can occur in membranes made from lipids with at least three moieties: carbon chains with low melting temperatures, chains with high melting temperatures, and a sterol (37–40). Accordingly, we used electroformation to produce giant unilamellar vesicles (GUVs) with three lipid components: DiPhyPC (*T*_melt_ < –120°C (41)), DPPC (*T*_melt_ = 41.3 ± 1.8°C (42)), and cholesterol (Figure 2, Supplemental Figure S2). DiPhyPC features branched (isopranyl) chains. These chains are also found in archaebacteria, where they may enhance survival in some extreme environments (34). A strong advantage of using DiPhyPC is that it lacks double bonds, so membrane photo-oxidation is minimized (44). Miscibility transition temperatures of GUV membranes of DiPhyPC/DPPC/cholesterol depend on the ratio of the three lipids (45). Above the transition, the lipids mix in one uniform phase, and below the transition they demix into liquid ordered (Lo) and liquid disordered (Ld) phases (45).

**Figure 2:**
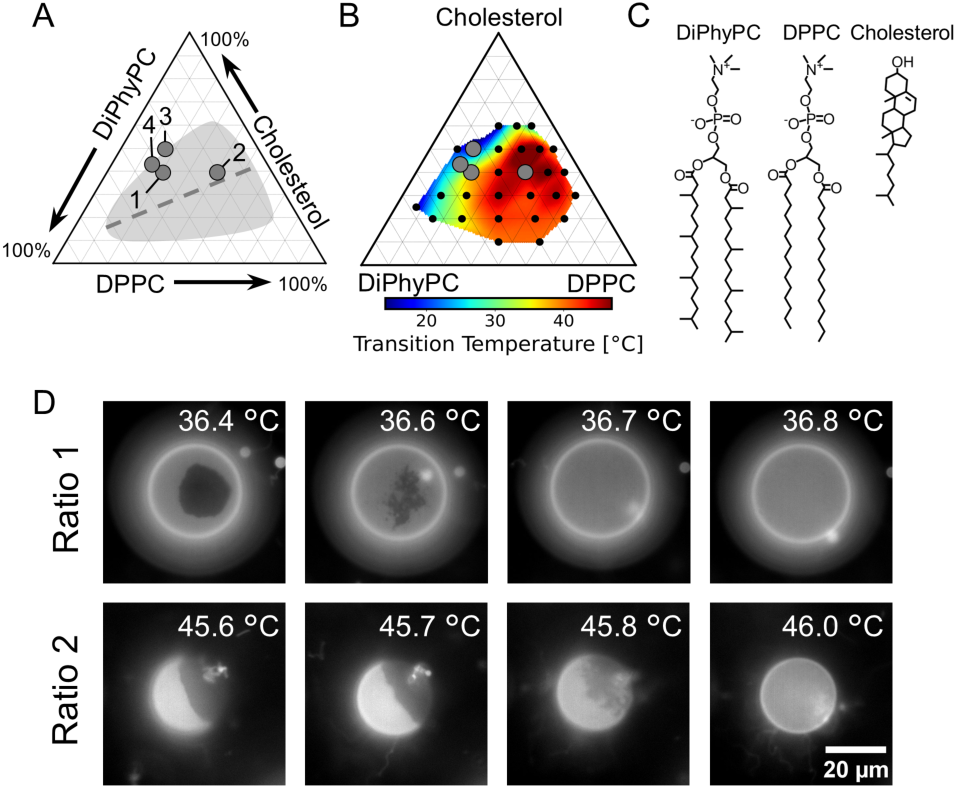
(A) Triangle representing all possible mixtures of three lipids: DiPhyPC, DPPC, and cholesterol. Vesicles were made from lipid ratios 1-4. The shaded area represents the approximate range of membrane compositions that demix into two liquid phases at 22 °C (35). The dashed line is a tie-line passing through 35/35/30 DiPhyPC/DPPC/cholesterol (35). (B) Map of the highest temperature at which two liquid phases are observed in vesicle membranes of DiPhyPC/DPPC/cholesterol (35). (C) Structures of DiPhyPC, DPPC, and cholesterol. (D) Fluorescence micrographs of giant unilamellar vesicles as a function of temperature, with 100 mM sucrose inside and outside the vesicles. Below the liquid-liquid transition temperature (36.6 °C for Ratio 1 and 45.9 °C for Ratio 2), the brighter phase is Ld, and the darker phase is Lo. Near the transition temperatures, fluctuations in domain shapes are characteristic of miscibility critical points (36). Above the transition, one uniform phase is observed. Images are representative and show additional structures in solution: smaller vesicles in images for Ratio 1 and bright lipid aggregates in images for Ratio 2.

### Single vesicles under osmotic pressure differences

The literature contains many examples of ternary vesicles being held above *T*_mix_, and then diluted into solutions of lower osmolarity. Often, this results in coexisting Lo and Ld domains appearing on the membranes, and then, if Δ*c* is large enough, vesicles cyclically rupturing (10, 11, 13, 25, 26). We confirm these results in Supplemental Movies S1 and S2. In isolation, these results imply that *T*_mix_ of the membrane always increases with tension. Occasionally, membrane domains appear and then disappear. It is tempting to interpret these events as an increase in *T*_mix_ followed by a decrease in *T*_mix_, but it is difficult to stringently rule out the possibility that local inhomogeneities in sucrose concentration caused the domains to disappear.

In Supplemental Figure S3 and Supplemental Movie S3, we observed a sequence of events that cannot be attributed to local sucrose inhomogeneities. When the exterior solution was diluted, the membrane of one vesicle exhibited three sequential states: domain formation, then domain disappearance, and, later, an abrupt decrease in vesicle radius, indicating rupture. Because rupture occurred without domains re-forming, the sequence can be interpreted as an increase in *T*_mix_ followed by a decrease in *T*_mix_. To find this single, rare vesicle, we conducted 40 independent experiments, each with ∼20 vesicles. The rarity of the event means the result should be regarded as preliminary. Another caveat is that in the final frames of the movie, it is not clear if the observed vesicle is touching another one.

Osmotic pressure differences are typically imposed on uniformly mixed membranes that are initially above *T*_mix_. We rounded out those studies by surveying vesicles that were initially slightly below *T*_mix_, so they started with coexisting Lo and Ld domains. When osmotic pressure differences were imposed, many of the vesicles ruptured. Over 20 independent experiments, each with ∼20 vesicles, none of the vesicle membranes became uniformly mixed. In other words, within this set of experiments, we did not find evidence of a decrease in *T*_mix_.

### Single vesicles under micropipette aspiration

To review, previous work has shown that when vesicles are pre-stressed into the stretching regime in which the area per lipid increases, applying more tension *decreases T*_mix_ of the membrane (11). Here, we find that when vesicles are not pre-stressed (such that significant thermal fluctuations are initially present), tension due to vesicle aspiration can *increase T*_mix_ of the membrane (Fig. 3). For example, Figure 3A shows a 3-component vesicle captured in a micropipette. When tension is applied at constant temperature, coexisting liquid domains appear in the membrane (Figure 3B and square symbol in Figure 3F). As tension is lowered, domains disappear (Figure 3C-E). We observed domains in five additional vesicles of a different lipid ratio (circle symbols in Figure 3F) at a constant temperature (20 °C, 25 °C, or 27 °C).

The broad spread in transition temperatures in Figure 3F is perplexing. It is similarly perplexing that three transitions in Figure 3F occur at tensions above 5 mN/m. For 1- and 2-component membranes, these higher tensions correspond to a membrane stretching regime, where the area per lipid increases (20). We presume (but do not definitively know) that the same cutoff applies to 3-component membranes. If so, then the high-tension transitions in Figure 3F contradict the assumption in the literature that tension always decreases *T*_mix_ in the stretching regime of area dilation (12). For the circles in Figure 3F, there is no clear correlation between the tension at the transition and either the sample temperature or the chronological order of the measurement (Supplemental Table S2). Scatter in transition tensions for this single sample may partly reflect vesicle-to-vesicle differences lipid compositions of ±2 mol% (as opposed to sample-to-sample differences, which are also estimated at ±2 mol%) because shifts in compositions translate into different *T*_mix_ values in tensionless membranes (35, 46–48).

At minimum, the observation that tension *increases T*_mix_ in Figure 3 means the previously reported *decrease* in *T*_mix_ is not characteristic of all aspiration experiments (11).

**Figure 3:**
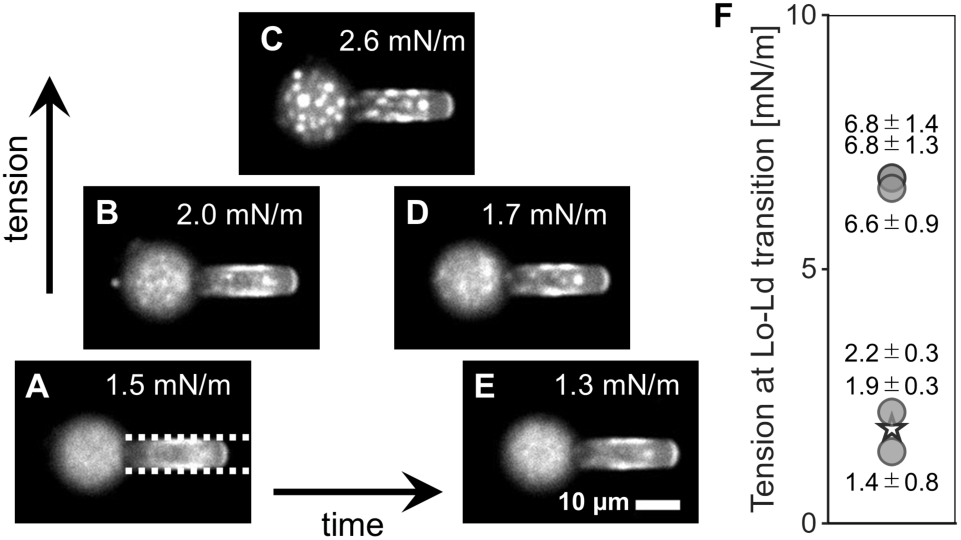
(A-E) Fluorescence micrographs from a timelapse movie at 28 °C of an aspirated vesicle with a composition corresponding to Ratio 3 in Figure 2A (Supplemental Movie S4). The main part of the vesicle is at the left, and a long “tongue” is pulled into the micropipette, which is shown by dashed lines. At low tensions, lipids in the main part of the vesicle (Panels A and E) mix uniformly. At intermediate tensions, fluctuations associated with critical phenomena are observed (Panels B and D). At high tension, coexisting liquid domains are observed (Panel C). (F) Star: Tension at which the membrane in Panels A-E demixes. Circles: Tensions at which membranes demix for five different aspirated vesicles made from a single electroformation batch of the lipid composition corresponding to Ratio 4 in Figure 2A. Two independent points overlap at 6.8 mN/m, one of which corresponds to Supplementary Movie S5. Temperatures of the vesicles range from 20 °C to 27 °C; experimental details are in Supplemental Table S2. Error bars represent the full range of tensions over which transitions could have occurred and arise predominantly from the image resolution, which limits the size of domains that can be identified.

### Populations of vesicles under osmotic pressure

Figures 4A and 4B illustrate three main qualitative results of population experiments in which vesicles are subjected to an osmolarity difference (Δ*c* = *c*_inside_ – *c*_outside_). First, as Δ*c* becomes negative such that vesicles deflate, *T*_mix_ remains approximately constant. In this regime, vesicles deflate as water transfers from the low-concentration sucrose solution inside the vesicles to the high-concentration solution outside. Every data point is normalized by plotting it as a shift (Δ*T*_mix_) in the miscibility temperature relative to *T*_mix_ of a population of vesicles from the same electroformation batch that were measured immediately after dilution into an isotonic solution. Second, as Δ*c* becomes positive and vesicles inflate, *T*_mix_ increases approximately linearly with the difference in osmolyte concentration, leading to positive values of Δ*T*_mix_. Third, at higher values of Δ*c*, a plateau is reached, possibly after a maximum.

**Figure 4:**
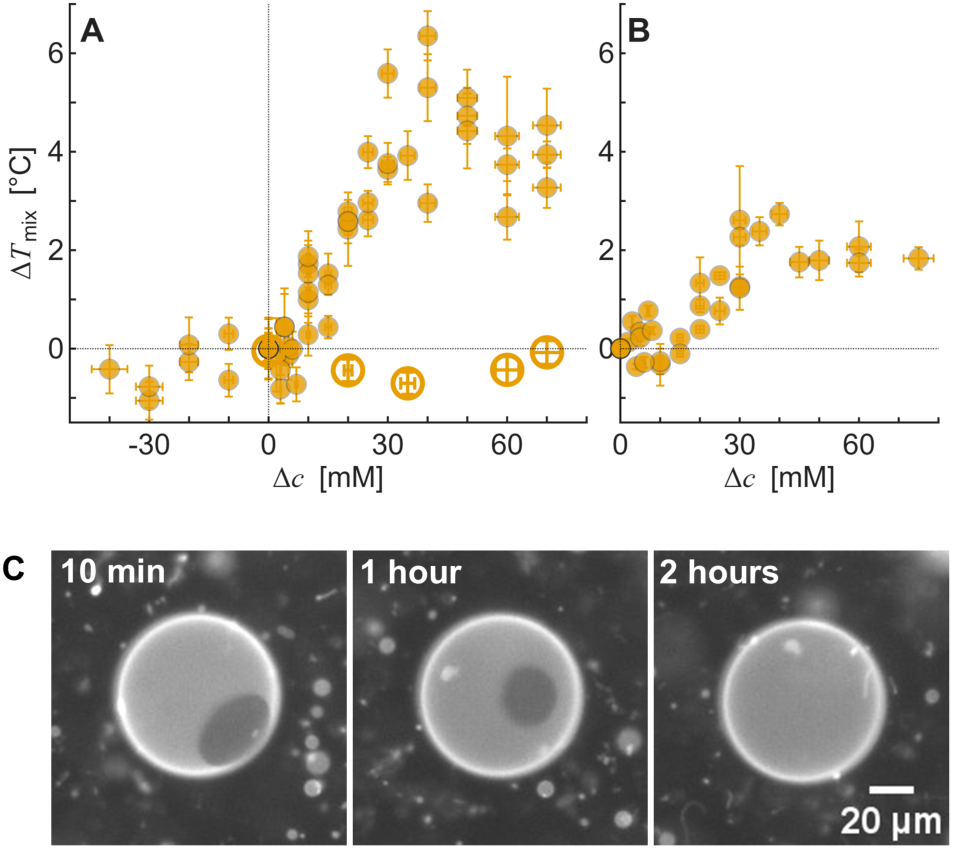
Shift in liquid-liquid phase transition temperatures (Δ*T*_mix_) for a population of GUVs initially in solutions of 100 mM sucrose, versus the difference in osmolarities inside and outside the vesicle (Δ*c* = *c*_inside_ – *c*_outside_). Filled circles denote data taken within 1 hour of applying the osmolarity difference, and each point represents a different experiment rather than an average. Open circles denote Δ*T*_mix_ values for samples measured 21 hours after dilution, including one at isotonic conditions. Δ*T*_mix_ of each data point is calculated relative to *T*_mix_ of vesicles that from the same batch and measured immediately after dilution into an isotonic solution, such that Δ*T*_mix_ = 0 is defined at Δ*c* = 0. Error bars are calculated as described in the Methods. (A) Vesicles of lipid ratio 1 from Figure 2A. Data are from 17 independent vesicle preparations. (B) Vesicles of lipid ratio 2 from Figure 2A. Data are from 6 independent vesicle preparations. (C) Time lapse of elimination of Lo-Ld coexistence in a single vesicle imaged at 38 °C at time points of 10 min, 1 hr, and 2 hrs after dilution to Δ*c* = 20 mM.

These three qualitative results hold for membranes with different ratios of the three lipids (Figure 4B and Supplemental Figures S4 and S5) and with a substitution of the saturated lipid DMPC in place of DPPC (Supplemental Figure S6). This substitution is noteworthy because the resulting membrane is the same as in previous micropipette experiments in which a decrease in *T*_mix_ was reported (11), namely 33/33/33 mol% DiPhyPC/DMPC/cholesterol. A similar decrease in transition temperature has been reported for binary mixtures with solid-liquid coexistence (10, 24).

The value of Δ*T*_mix_ at the plateau depends on the membrane’s lipid composition. The area fraction of dark, Lo phase increases from Panel A to Panel B in Figure 4 (corresponding to a change from Ratio 1 to Ratio 2 in Figure 2A), and the plateau value decreases. Other researchers have found a larger Δ*T*_mix_ for liquid-liquid transitions than solid-liquid transitions (24) and a nonmonotonic dependence of Δ*T*_mix_ on the area fraction of solid phase (10).

To minimize errors in Figure 4A and 4B, we split every vesicle solution into control and test samples. We diluted the control into an isotonic solution of the same osmolyte, and we diluted the test sample into a solution of lower (positive Δ*c*) or higher (negative Δ*c*) osmolyte concentration. This procedure addresses the problem that electroformation (and other methods of forming vesicles) result in small, day-to-day variations in lipid ratio (37).

Several challenges arise when attempting to quantitatively evaluate *T*_mix_ as a function of osmolarity differences. It is easy to misinterpret values from the slope and plateau regions of Figure 4. At low osmotic pressures, Δ*T*_mix_ is approximately fit by a line that passes through the origin. A compelling (but incorrect) conclusion is that none of the vesicles have hidden area in tubes or lipid aggregates. Similarly, the onset of the plateau is at high osmotic pressures. A compelling (but incorrect) conclusion is that it represents the start of vesicle lysis. As we will see below, membrane tension cannot be directly inferred from Δ*c* due to two significant effects: the presence of large reservoirs of hidden area and the occurrence of membrane ruptures. These two effects also present challenges in considering Δ*c* as an independent variable.

There are four lines of evidence that freshly-made GUVs have significant hidden area. First, tubes retract into GUVs as osmotic pressure is applied (18), as shown in Figure 5A. Second, most GUVs have no measurable tension (33), even under an osmolarity difference of +20 mM (Figure 5B). At higher osmolarity differences (Δ*c* = +40 mM) membrane tension can still be zero. Third, the radii of individual GUVs increase dramatically before vesicles rupture, corresponding to an ersatz osmolarity change as high as ∼30 mM (Figure 5C). The ersatz change in internal concentration Δ*c* is calculated as Δ*c/c*_0_ = (1 – (1 + Δ*r*/100)^-3^) where *c*_0_ is the initial internal sucrose concentration (100 mM) and Δ*r* is the percent change in radius. This equation is simply a statement that the membrane is permeable to only water (and not sucrose) and that the final sucrose concentrations inside and outside the vesicle are equal. Fourth, positive osmotic pressure increases the width of the sigmoidal curves (an indicator of the variation) used to determine *T*_mix_ of a vesicle population (Supplemental Figure S7).

**Figure 5:**
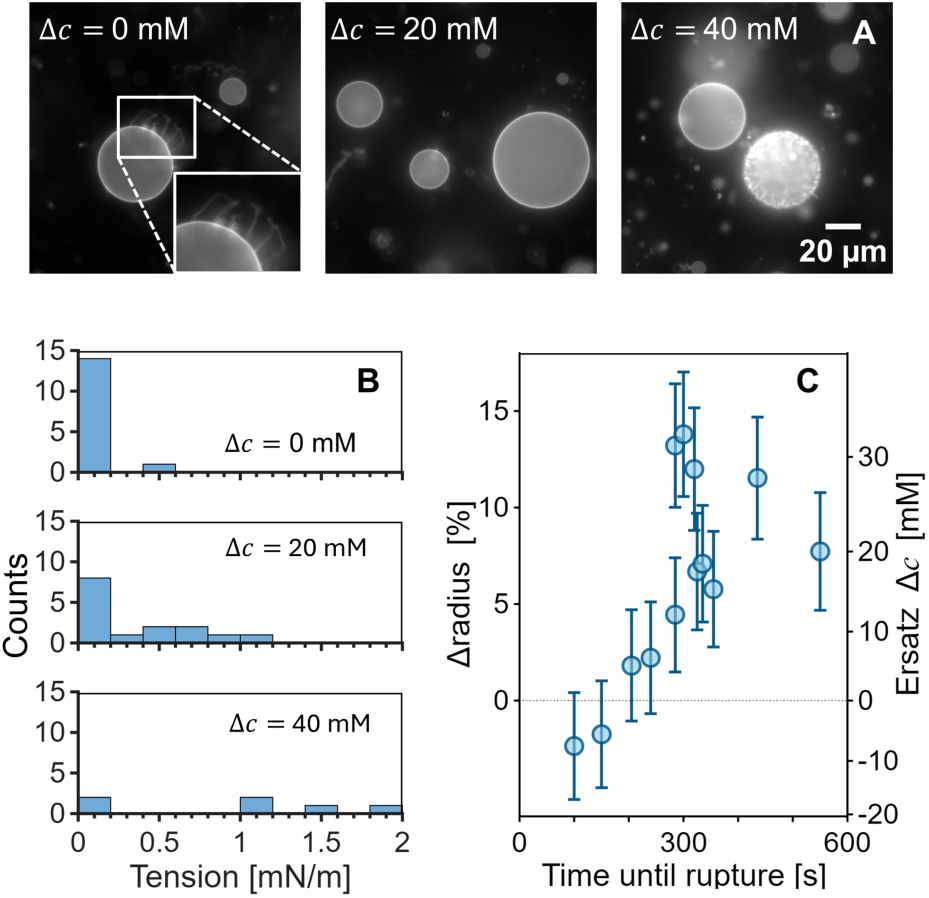
Evidence of hidden area in vesicles. The lipid composition corresponds to point 1 in Figure 2A. (A) Retraction of tubules and lipid aggregates as the osmotic pressure difference increases. At Δ*c* = 0 mM, a bright halo of tubules and some aggregates are visible; the inset shows higher magnification. At Δ*c* = 20 mM, fewer tubules and aggregates are visible. By Δ*c* = 40 mM, many vesicles have ruptured, which can cause dense vesiculation, tubulation, and aggregate formation. (B) Membrane tension of individual vesicles measured by micropipette aspiration for osmotic pressure differences of Δ*c* = 0, 20, and 40 mM. At Δ*c* = 20 mM, roughly half of vesicles have tensions near 0 mN/m. Even at the highest osmolarity difference of 40 mM, some vesicles still have tensions near 0 mN/m. At 40 mM, tension could be measured for only about half of vesicles because some vesicles had previously ruptured and had very low tensions (so they were drawn completely into the pipette) and others had very high tension (so they ruptured upon application of more tension). Consequently, vesicles at the lowest and highest tensions are likely underrepresented in the data. Data are tabulated in Supplemental Table S3. Among all vesicles prepared with lipid ratio 1, only those in this experiment used 0.8 mol% of the fluorescent dye Rhod-DOPE rather than Rhod-DPPE. (C) Data from a time lapse of a single population of vesicles (Supplemental Movie S1). Under an osmotic pressure difference, vesicles swell and eventually burst. Vesicles that persist the longest before rupture have the largest change in radius, corresponding to ersatz osmolarity changes as high as ∼30 mM. Error bars for Δradius reflect a 2% uncertainty in the initial radius and the radius immediately before rupture. Uncertainty arises from image pixelation: measurements at different angles through the center of a vesicle result in different apparent radii. Data are tabulated in Supplemental Table S4.

Hidden area allows vesicles to persist at higher osmolarity differences. We can calculate the value of Δ*c* at which all vesicles would rupture if they had no hidden area. If a spherical vesicle lyses when its surface area increases by ≤ 4% (21, 23), then for a vesicle with an initial radius of 10.0 µm, a 4% increase in surface area corresponds to a final radius of only 10.2 µm. Put another way, if vesicles that were electroformed in a 100 mM solution had no hidden area, we would expect them to lyse at an osmolarity difference of only Δ*c* = +5.8 mM. In contrast, applying Δ*c* = 40 mM to a spherical vesicle with an initial radius of 10.0 µm (by diluting the solution from 100 mM to 60 mM) corresponds to a much larger final radius of 11.9 µm. This quantitative inconsistency cannot be due to slow permeability of water across vesicle membranes. Osmotic pressures equilibrate across membranes of GUVs in roughly 5 min. (23, 38), an order of magnitude faster than the timescale of our experiments.

The most convincing evidence that ruptures occur before the plateau region is visual: some vesicle radii abruptly decrease at values of Δ*c* well before the plateau (∼30 mM, Supplemental Movie S6). Ruptures cause flow of solution from the inside to the outside of vesicles, as previously seen by expulsion of encapsulated aggregates (22), vesicles (16), and dye (13).

Techniques that test for flow in the opposite direction, for example by sensing the movement of dye from the outside of vesicles to the inside, are less effective (10). Supplemental Movie S7 and Supplemental Figure S8 show that rupture of osmotically inflated vesicles is not necessarily accompanied by a measurable influx of calcein from the external solution. Observation of pores in only some, but not all, vesicles at low values of Δ*c* is best explained by a wide range of hidden area (Figure 5B), not by the range of swelling rates (Supplemental Table S1). Evidence that pores, whether large or small, occur in all vesicles is that Δ*T*_mix_ drops to zero over long experimental times (e.g., 21 hours for the open circles in Figure 4A), accompanied by the visual elimination of Lo-Ld coexistence that returns the vesicle to its original mixed state (Figure 4C).

Another challenge in quantitatively assessing tension in osmotic pressure experiments is that an increase in temperature causes a decrease in membrane tension. When osmotic pressure causes an increase in *T*_mix_, experimental temperatures must be raised to measure the new *T*_mix_ value, which results in a decrease in tension. Tension decreases 0.34 mN/m per °C for a disordered membrane of eggPC lipids (21), and about twice as much for a more ordered membrane of DMPC (because the thermal area expansivity of DMPC is about twice as large as eggPC) (39). Thermal expansions due to melting of solid phases are an order of magnitude larger (39), which may help explain why shifts in membrane melting temperatures measured in osmotic pressure experiments are smaller than shifts in *T*_mix_ of liquid phases (24). Chen and Santore used this effect to intentionally change membrane tension, with the caveat that the technique has large uncertainties (31).

To summarize, the presence of significant hidden area presents major challenges in considering Δ*c* as a quantitative, independent variable. Moreover, when vesicles have pores and/or rupture, only an apparent Δ*c* can be stated for a population of vesicles, and the corresponding tension varies with temperature. Although it is tempting to use the Laplace equation to directly convert Δ*c* values into membrane tensions for a vesicle population, it is not clear if the resulting tensions are meaningful beyond setting bounds. For example, the onset of the plateau region can give an upper bound on the amount of hidden area in newly formed vesicles. A separate challenge arises from the Laplace equation: larger vesicles are subject to bigger tensions at a given Δ*c.* Different techniques produce vesicles of different sizes, and, presumably, different excess areas. Experimentally, electroformed vesicles have nonuniform sizes that increase with preparation time (40), which complicates quantitative comparisons between laboratories, even if they all use electroformation.

To our knowledge, only one technique definitively translates osmotic pressures on single vesicles into membrane tensions (41). However, the technique requires analysis of rates of pore formation on individual GUVs that lack hidden area, are subjected only to small Δ*c* (e.g., 4 mM), and are aspirated in a micropipette as osmotic pressure differences are applied (41). In principle, such a measurement would have to be repeated for every new lipid composition, due to varying edge tensions and lysis tensions (42). Nevertheless, this technique has shown that membrane tension does increase in response to osmotic pressures low enough to avoid poration (41).

### Other concentrations, osmolytes, and types of lipids

Challenges in using the variable Δ*c* extend beyond the problem of hidden area, pores, and ruptures. Osmotic pressure differences are accommodated by changes in the volumes of vesicles. Therefore, Δ*c* should be rescaled so that the volume change is accounted for. For positive values of Δ*c*, when the concentration of osmolyte is initially higher inside the vesicle than outside, the two osmolyte concentrations will be equal after the vesicle gains a volume of water equivalent to Δ*c*/*c*_outside_ of its initial volume.

The benefit of rescaling is clear in Figure 6. Panels 6A and 6B show the shift in miscibility temperatures versus Δ*c* in vesicles with different initial osmolyte concentrations (100 mM, 150 mM, and 200 mM) diluted into solutions with lower osmolarity. Rescaling the *x*-axis of each data set by a factor of Δ*c*/*c*_outside_ overlays the data sets (Figure 6C and 6D). In the rescaled data, Δ*T*_mix_ increases with Δ*c* /*c*_outside_ up to a value of ∼0.5, corresponding to ∼50% increase in volume and ∼30% increase in apparent surface area, consistent with evidence for significant hidden area in Figure 5 (with raw data in Supplemental Figures S4, S5, S9, S10, and S11). Although rescaling mitigates scatter in data due to different initial osmolarities, it does not mitigate problems due to hidden area, pores, and ruptures.

**Figure 6:**
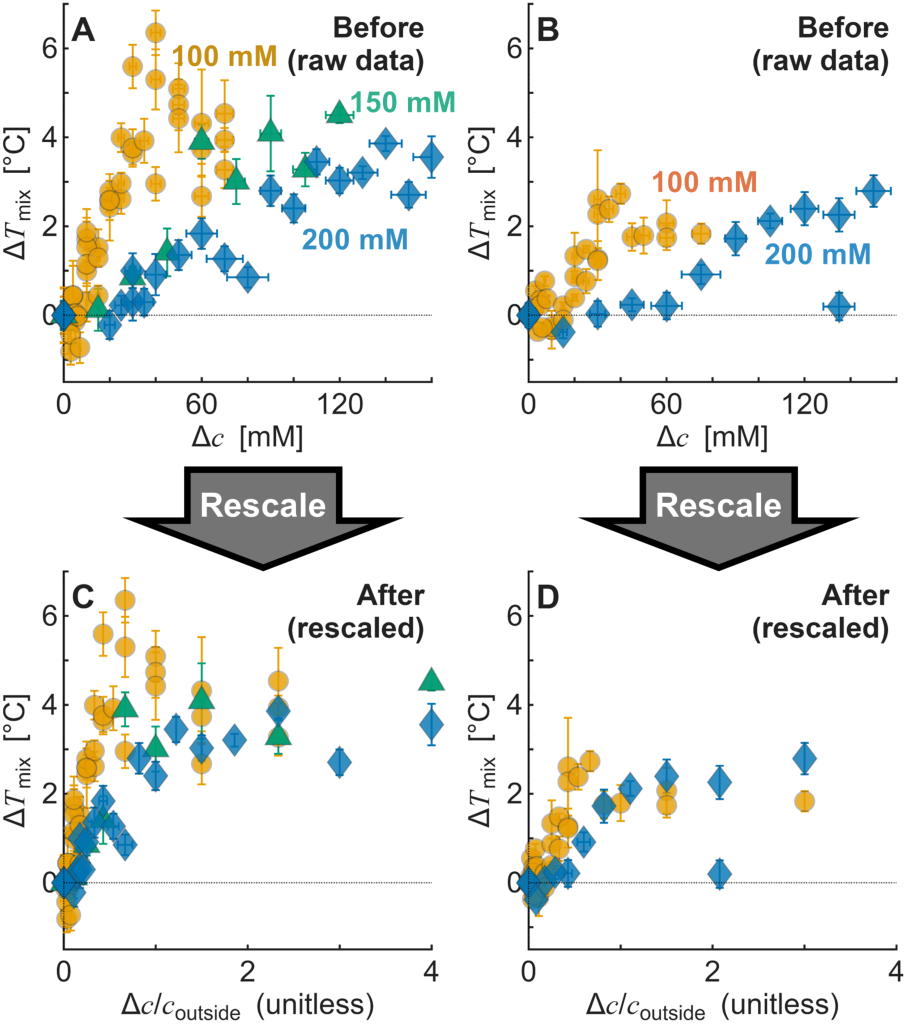
(A,B) Raw data for shifts in liquid-liquid phase transition temperatures of GUV membranes with initial sucrose concentrations inside and outside the vesicles of 100 mM (circles, repeated from Figure 2), 150 mM (triangles), and 200 mM (diamonds). Data for 100 mM sucrose are repeated from Figure 2. Data in Panel A is for vesicles of composition 1, and data in Panel B is for composition 2. Data are from 23 and 8 independent vesicle preparations in Panels A and B, respectively. Error bars are calculated as described in the Methods. (C,D) Rescaled data from Panels A and B with the x-axis re-plotted as the relative concentration difference, Δ*c*/*c*_outside_.

Another experimental condition that changes *T*_mix_ is the choice of molecule used as an osmolyte. Although osmolytes are often modeled as ideal solutes with only colligative effects, many osmolytes interact with membrane headgroups and/or create depletion layers (43). The experiments in Figures 2-6 were conducted with sucrose solutions. Symmetric application of sucrose across uncharged membranes does not shift *T*_mix_ in vesicles (44).

Repeating the experiments in Figure 4A with TMAO instead of sucrose resulted in the same qualitative behavior, with lower shifts in *T*_mix_ (Supplemental Figures S12, S13, S14). Other researchers have found that even symmetric solutions of dextran (40 kD and 200 kD) and PEG (6 kD) shift *T*_mix_ (25). Asymmetric concentrations of these molecules (and of glucose) increase *T*_mix_ (25). Similarly, when different types of salt solutions are applied across a charged membrane, *T*_mix_ shifts (45). Even simple sugars can generate small membrane tensions when different sugars are applied inside vs. outside vesicles (∼10^−8^ to 10^−5^ mN/m) (46, 47).

A third experimental condition of interest is the lipid composition of the membrane. Figure 6A and 6B show that changing the ratio of the lipids affects the magnitude of Δ*T*_mix_ when a difference in osmotic pressure is imposed. Similarly, Wongsirojkul et al. changed the type of lipids (using DOPC rather than DiPhyPC in ternary membranes) and, depending on the lipid ratio, found Δ*T*_mix_ plateau values of ∼12-16 °C, roughly twice the values we measure (24).

Changing a membrane’s lipid composition also affects the bending stiffness of each phase. The bending stiffness of a uniformly mixed membrane is typically lower than the weighted average of the Lo and Ld phases (Supplementary Figure S17). Based on hints in the literature (48, 49), we investigated membranes that had the potential to exhibit the opposite behavior. Charged lipids have been reported to have a non-linear stiffening effect on membranes (48), so we investigated vesicles with 10 mol% charged DPPS (plus 6% DiPhyPC, 44% DPPC, and 40% cholesterol). We did not observe a decrease in *T*_mix_ (nor a significant increase) when we imposed an osmotic pressure difference across these vesicles (Supplemental Figure S15). Other lipid compositions produce Lo and Ld phases with more equal thicknesses (rather than a much thicker Lo phase) (49). For one such composition (6/54/40 mol% DiPhyPC/Di(13:0)PC/ cholesterol, with 0.8 mol% Rhodamine DPPE), we did not observe a decrease in *T*_mix_ (Supplemental Figure S15).

## DISCUSSION

Tension has been predicted to increase *T*_mix_ or to decrease *T*_mix,_ depending on whether tension predominantly suppresses out-of-plane thermal fluctuations of the membrane (50), as is asserted to occur at low tensions (19), or if tension increases the lateral area per lipid (15, 31), as is asserted to occur at higher tensions (19). To help build intuition, we will briefly summarize each approach.

Thermal fluctuations cause membrane bending. Gordon et al. proposed a model in which demixing of a membrane into equal fractions of a disordered, A-rich phase and an ordered, B-rich phase results in a change in free energy per area proportional to *ln*[(*κ*_A_*κ*_B_) / *κ*_αΒ_^2^], where *κ* is the bending stiffness ((50) and references within). They note that the bending stiffness of the Lo phase is significantly larger than that of the Ld phase (K_Lo_ > 2K_Ld_) and that the bending stiffness of the mixed state is close to that of the Ld phase (K_Ld_ ∼ K_mixed_). As a result, fluctuations in a phase-separated membrane incur a free energy cost (a positive number proportional to ln2), so mixing is favored. Conversely, suppressing fluctuations favors phase separation. Therefore, in this approach, tension increases *T*_mix_. For arbitrary fractions of A and B components, Markin considers summing the inverses of curvature elasticities (51).

Approaches based on thermal fluctuations are compelling for explaining results from two types of experiments: the osmotic pressure experiments in Figure 4, Figure 6, and many previous papers (9, 10, 12, 24, 25), and the micropipette aspiration experiments in Figure 3. For solid and liquid transitions in membranes, Gordon et al. predict Δ*T*_mix_ of about +3K (50). In the fluctuation approach, the largest values of Δ*T*_mix_ should occur in membranes for which the bending stiffness of the mixed membrane differs most from the bending stiffness of the weighted average of the Lo and Ld phases (Supplemental Figure S17), which is presumably near the middle of a tie-line rather than an edge. We are not aware of any membrane that should have the opposite effect, although we explored potential candidate membranes containing charged lipids and alternative chain lengths (Supplemental Figure S15).

To build intuition for the second approach, it is useful to consider a block of aluminum. At high temperature, aluminum is a liquid, which has a lower density than the solid. From the Clausius-Clapeyron equation, pressure drives systems toward higher-density states. Therefore, negative pressure favors the liquid and decreases the melting temperature. Membranes are a 2-dimensional analogy of the block of aluminum. The high-temperature state of a ternary membrane has a lower areal density than the weighted average of the Lo and Ld phases (Supplemental Figure S17). Tension drives the system to the low-density, mixed state. Therefore, in this approach, tension decreases *T*_mix_. Chen et al. used this thermodynamic approach to quantitatively explain the decrease in the solid-liquid transition of a bilayer under tension (31). In an earlier model, Uline et al. considered stretching in context of a Gibbs-Duhem equation, with molecular area and tension as conjugate variables (15).

This second approach is compelling in explaining decreases in *T*_mix_ in experiments that isolate stretching using supported bilayers (26), aspirated vesicles (11, 31), and vesicles under hydrostatic pressure (27–30). The model of Uline et al. predicts a decrease in *T*_mix_ that quantitatively agrees with experimental results by Portet et al., who used micropipette aspiration to bypass the fluctuation regime and access the stretching regime (11). However, this second approach does not explain why the onset of phase separation in some membranes is at tensions higher than 5 mN/m, which is presumably in the stretching regime (Figure 3F).

Given that some of the experimental results are consistent with tension *increasing T*_mix_ and others are consistent with tension *decreasing T*_mix_, it is worthwhile to explore whether any approach might accommodate both types of results. Here, we introduce two speculative scenarios in which tension causes nonmonotonic changes in *T*_mix_ (Fig. 7), as a possible interpretation of the combined evidence. Below, we show that although both scenarios have advantages, neither scenario completely reconciles the contradictory experimental results.

Both scenarios start with an initial condition in which vesicles have hidden area, which has no measurable effect on *T*_mix_. The scenarios differ in their intermediate states. In one scenario, a small decrease is followed by a large increase (a shape colloquially termed a “checkmark”). In the other scenario, the order is reversed (an “inverse checkmark”). The scenarios are presented for single vesicles (on the left of Figure 7) and for populations of vesicles (on the right of Figure 7). Once a pore opens in a vesicle, as observed when osmotic pressure is applied, only an apparent Δ*c* is measured (Fig. 1). Both scenarios end with a final condition in which vesicles rupture, and *T*_mix_ decreases sharply.

**Figure 7:**
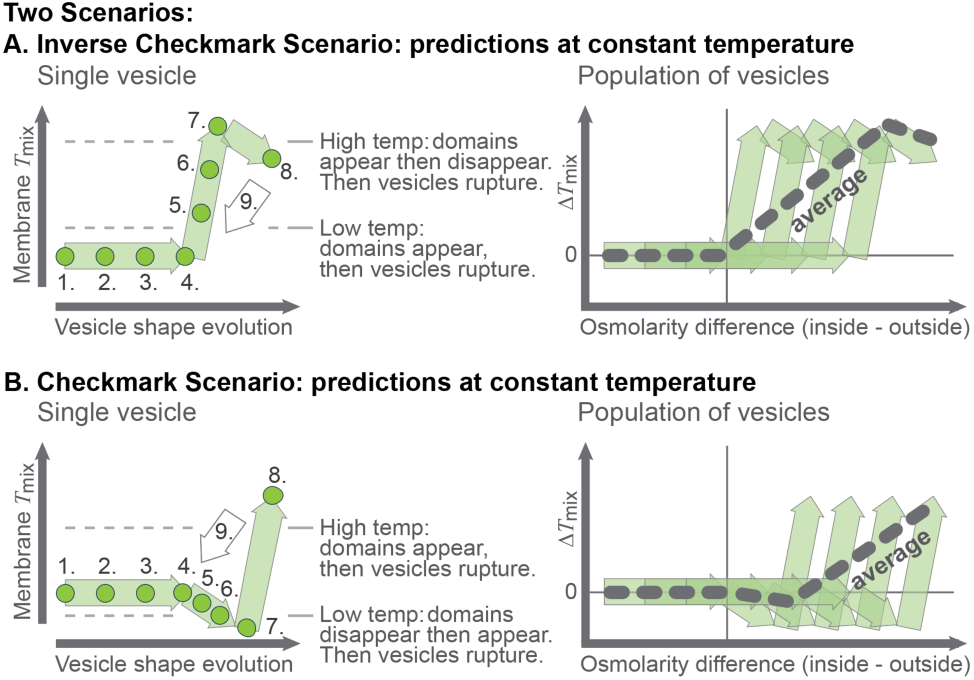
(A) Predictions from the Inverse Checkmark Scenario for experiments with single vesicles (left) and vesicle populations (right). Numbers correspond to the vesicle states in Figure 1. (B) Predictions from the Checkmark Scenario.

### Inverse Checkmark Scenario

We will first describe how predictions of the Inverse Checkmark Scenario are either consistent with current results, are inconsistent, or are missing from the experimental record. The Inverse Checkmark Scenario spans four regimes:

- First, when hidden area retracts, no change in *T*_mix_ (in other words, Δ*T*_mix_= 0). Roughly half of newly electroformed vesicles have no measurable tension (Figure 5). Although new vesicles are typically taut enough that domains merge (52), the hidden area of a vesicle can be so large that some vesicles with Δ*c* ≥ 20 mM have zero tension. As a result, quantitative evaluation of tensions from Δ*c* values are nearly impossible. Plasma membranes of cells also have reservoirs of excess area that mitigate tension (53).
- Second, a large increase in Δ*T*_mix_ occurs, as in points 4-7 in Figure 7A. Low tensions are asserted to predominantly suppress out-of-plane thermal fluctuations of the membrane (19). Corresponding theoretical predictions are that Δ*T*_mix_ should increase (50). Indeed, an increase in Δ*T*_mix_ is experimentally observed when osmotic pressure is applied to either single vesicles or a population of vesicles (Figures 4 and 6 and references (9, 10, 12, 24, 25)) or when micropipette aspiration is applied to single vesicles at low tensions (Figure 3). These aspiration results at low tensions are important because they show that an increase in Δ*T*_mix_ is not limited to experiments that impose osmotic pressure differences.
- Third, a small decrease in Δ*T*_mix_ occurs, as in points 7-8 of Figure 7A. Higher tensions are asserted to predominantly increase the area per lipid (19). Corresponding theoretical predictions are that Δ*T*_mix_ should decrease (15, 31). Indeed, a decrease in Δ*T*_mix_ is experimentally observed when micropipette aspiration is applied to single vesicles that are pre-set in the stretching regime (11). Similarly, tension decreases melting temperatures of vesicle membranes (31). Likewise, Illouz et al. isolated the effects of stretching from effects of fluctuations by stretching flat, supported lipid bilayers and saw a decrease in Δ*T*_mix_ (26). The same conclusion was reached when hydrostatic pressure was applied to vesicles, which causes lateral compression of membranes (figure 4a of (30)). Because pressure causes an increase in *T*_mix_ (27–29), the results are consistent with membrane stretching causing a decrease in *T*_mix_.

Inverse Checkmark Scenario also makes a prediction that spans the second and third states. Figure 7A shows a horizontal line at a constant, high temperature as tension (or Δ*c*) is applied. To unambiguously observe an increase in Δ*T*_mix_ followed by a decrease Δ*T*_mix_, the experimental temperature must fortuitously lie below and very close to the highest possible *T*_mix_ for whichever vesicle is being observed. In our experiments, we set an even higher criterion. To rule out any effects of local inhomogeneities in sucrose concentration, we searched for vesicles that exhibited three sequential steps: domain formation, domain disappearance, and, later, an abrupt decrease in vesicle radius. We observed this sequence in a single osmotically stressed vesicle, out of a set of ∼800 vesicles (Supplemental Figure S3). However, that single case is not ideal; we cannot tell if the vesicle exchanges lipids with another vesicle at the end of the movie (though if it were, we would not expect it to rupture nor for its radius to decrease). Given that we observed only one event, this result should be considered preliminary.

One of our key experimental observations appears inconsistent with a decrease in Δ*T*_mix_ in the stretching regime: when single vesicles are aspirated at a constant temperature, domains sometimes appear at tensions > 5 mN/m, which are high enough to presumably be in the stretching regime (Figure 3). If so, then our data are inconsistent with *T*_mix_ only decreasing in the stretching regime.

- Fourth, membrane tension Δ*T*_mix_ sharply decreases due to small pores or large ruptures (as in Supplemental Figure S3). Given that Δ*T*_mix_ decays to zero over time (Figure 4A), any imposed Δ*c* should be treated as an apparent, time-dependent osmolarity difference.

In conclusion, many (but not all) of the results in this manuscript and in the literature are consistent with the Inverse Checkmark Scenario.

### Checkmark Scenario

Next, we will describe how predictions of the Checkmark Scenario are either consistent with current results, are inconsistent, or are missing from the experimental record over the four regimes:

- First, when hidden area retracts, Δ*T*_mix_= 0. This regime is the same as in the Inverse Checkmark Scenario above.
- Second, a small decrease in Δ*T*_mix_ occurs, as in points 4-7 in Figure 7A. Any support of this prediction comes with significant caveats. For example, although the equations used to predict a decrease in Δ*T*_mix_ are most relevant over small changes (15, 31), they are applied to increases in the area per lipid, which are asserted to dominate at higher tensions (19). Likewise, although decreases in Δ*T*_mix_ are observed experimentally, they are restricted to cases in which the area per lipid increases (10, 26–29, 31).
- Third, a large increase in Δ*T*_mix_ occurs, as in points 7-8 in Figure 7A. The strongest support of this prediction comes from the higher transition tensions in Figure 3F. However, although a decrease in Δ*T*_mix_ has been theoretically predicted, it applies to suppression of fluctuations (50), which are asserted to dominate at lower tensions (19).

The Checkmark Scenario also makes a prediction that spans the second and third states. Figure 7B shows a horizontal line at a constant, low temperature as tension (or Δ*c*) is applied at constant temperature. The prediction is difficult to validate in an experiment at constant temperature. To observe an unambiguous *decrease* in *T*_mix_, the experimental temperature must fortuitously lie below and very close to *T*_mix_ of a tensionless vesicle. As tension is applied, coexisting liquid domains should disappear due to a decrease in *T*_mix_, then reappear due to an increase in *T*_mix_. We were unable to find a single vesicle that met these criteria within 20 independent experiments, each with ∼20 vesicles.

- Fourth, membrane tension Δ*T*_mix_ sharply decreases due to small pores or large ruptures. This regime is the same as in the Inverse Checkmark Scenario.

In conclusion, the Checkmark Scenario is in better quantitative agreement with the measured aspiration tensions in Figure 3. However, no other experimental data strongly support the scenario, and it is difficult to theoretically justify why a decrease in *T*_mix_ would precede an increase in *T*_mix_. We have no data from individual, osmotically stressed vesicles in support of the nonmonotonic behavior of the Checkmark Scenario.

### Summary

Overall, our discussion of the two scenarios shows that enormous opportunities remain for the development of theory and simulations that quantitatively predict how tension increases or decreases *T*_mix_ under different experimental conditions. On the experimental side, our results suggest a roadmap of future tests that could assess whether *T*_mix_ varies with tension in a nonmonotonic way (as in the Inverse Checkmark and Checkmark scenarios) or in a purely monotonic way. An ideal experiment would use micropipette aspiration with dynamic temperature control (11), which we did not have access to in our experiments. To sequentially explore regimes in which fluctuations decrease and the area per lipid increases, vesicle should initially have as little tension as possible, rather than being pre-stressed into the stretching regime. We also recommend ample patience and funding, because experience (11) tells us that these experiments will likely be as challenging as they are rewarding.

Independent of whether either scenario is correct, the data in Figure 4 can help researchers determine if significant errors in *T*_mix_ arise from experimental conditions that affect vesicle tension. For example, the low and high viscosity osmotic conditions in Stanich et al. correspond to Δ*c*/*c*_outside_ of ∼40 and ∼0.04, respectively (54). If those experiments had measured *T*_mix_, the first condition would have caused a systematic shift in *T*_mix_ as large as the plateau value, and the second would have caused no measurable shift. Likewise, osmotic shifts explain why more liquid-liquid phase separation was observed in binary membranes (made of PChemsPC and diPhyPC) at Δ*c*/*c*_outside_ = 0.25 than at 0.0 (55). Another condition that may affect membrane tension is addition or depletion of lipids or sterols via cyclodextrin (56, 57). Our observation that most new GUVs are tensionless and contain hidden area mitigates concerns that small additions or depletions of lipids shift *T*_mix_, although severe depletion may still shift *T*_mix_. Another way to impose tension is through intentional decreases in temperature (31). In our experiments, vesicles were cooled from electroformation temperatures of 55-60 °C down to 20 °C and remained largely tensionless. However, if we had used lipids with higher melting temperatures, we would have needed higher electroformation temperatures (32). Researchers may also introduce errors simply by changing temperature to measure *T*_mix_, because thermal cycling decreases membrane tension and because some procedures cause hysteresis in *T*_mix_ (Supplemental Figure S16).

## CONCLUSIONS

To trigger liquid-liquid phase separation, researchers have historically varied molecular characteristics of a membrane (such as the types, ratios, and crosslinking of membrane components) as well as its physical parameters (such as temperature, hydrostatic pressure, and tension). Here, we have investigated two techniques in which membrane tension can increase the liquid-liquid transition temperature of a membrane: micropipette aspiration and differences in solution osmolarities. When newly formed vesicles are aspirated at constant temperature, the tension causes membranes to phase separate. Quantitative values of vesicle tension are straightforward to extract from aspiration experiments. Swelling of vesicles via differences in osmolyte concentrations also causes membranes to phase separate. However, electroformed vesicles have a wide distribution of hidden areas, which means that differences in solvent concentrations, Δ*c*, are difficult to convert into tensions. Moreover, vesicles rupture over a wide range of Δ*c* values, so Δ*c* should be viewed as only an apparent value. Challenges in comparing between experiments with different osmotic conditions is partially mitigated by reporting the relative change in concentration, Δ*c*/*c*_outside_ rather than Δ*c*.

For the theory and simulation communities, our results provide an opportunity to reconcile models written to explain an increase in *T*_mix_ due to decreases in out-of-plane fluctuations (50) and a decrease in *T*_mix_ due to membrane stretching (15, 31). For the experimental community, our results suggest experiments that span regimes of low tension and high tension, to test whether *T*_mix_ varies monotonically with tension or if it varies non-monotonically as in the scenarios we present. In a broader biological context, tension in a cell’s plasma membrane affects several inter-related characteristics, including contractility, motility, morphology, cytoskeletal networks, exocytosis, and vesicle trafficking (53). When tension couples to liquid-liquid phase separation, signals from channels and proteins that partition differently into each phase can potentially be amplified.

## ACKNOWLEDGMENTS

S.L.K. acknowledges funding from NSF MCB-2325819. D.A.F. acknowledges support from the NSF Center for Cellular Construction (DBI-1548297) and a Chan Zuckerberg Biohub Investigator Award. We thank Lucia Parolini and Pietro Cicuta for exploratory aspiration experiments in 2016.

## DATA AND CODE AVAILABILITY STATEMENT

All data used in constructing all plots are available in the Supporting Material. Code to calculate *T*_mix_ was implemented in two different (and functionally equivalent) ways. Code to calculate *T*_mix_ in Supplemental Figure S15 is available on GitHub at https://github.com/kentwils/mixing_temp_fit. Code to calculate *T*_mix_ for Figures 4, 6 and Supplemental Figures S4, S5, S6, S7, S9, S11, S12, S13 and S14 is available on GitHub at https://github.com/tkuroy/guv_tmix_analysis, using data in Figure 4, Figure 6, and Supplemental Figure S12 as an example.

## DECLARATION OF GENERATIVE AI AND AI-ASSISTED TECHNOLOGIES IN THE MANUSCRIPT PREPARATION PROCESS

ChatGPT (OpenAI; models GPT-5 and GPT-6) was used to revise existing code for calculating *T*_mix_ (available on GitHub at https://github.com/kentwils/mixing_temp_fit) to simplify calculations of *T*_mix_ in Figures 4, 6 and Supplemental Figures S4, S5, S6, S7, S9, S11, S12, S13 and S14 (available on GitHub at https://github.com/tkuroy/guv_tmix_analysis).

## AUTHOR CONTRIBUTIONS

T.K., K.J.W., A.C., and C.E.C. performed research. T.K. and K.J.W. analyzed data. T.K., K.J.W., S.L.K. designed research and wrote the manuscript. C.E.C., A.C., and D.A.F. provided feedback on the manuscript.

## DECLARATION OF INTERESTS

The authors declare no competing interests.

## Notes

### Competing Interest Statement

The authors have declared no competing interest.

### Summary of Updates

The text has been updated throughout based on feedback to be more clear. Most sections have been revised, especially the discussion section, which includes a clearer and more detailed explanation of the "Checkmark" and "Inverse Checkmark" scenarios.

