## Supplemental Information for "Ups and downs of liquid-liquid transitions in GUV membranes from osmolarity and aspiration tensions"

**Contents:**

– Supplemental Figures S1-S17

– Supplemental Table S1-S4

**Supplemental Figures:**

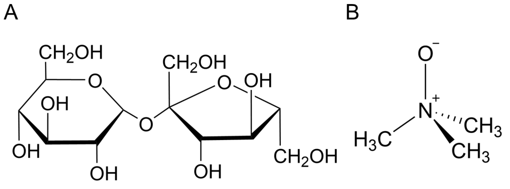

**Figure S1:** Chemical structures of A) sucrose, B) TMAO (trimethylamine *N*-oxide)

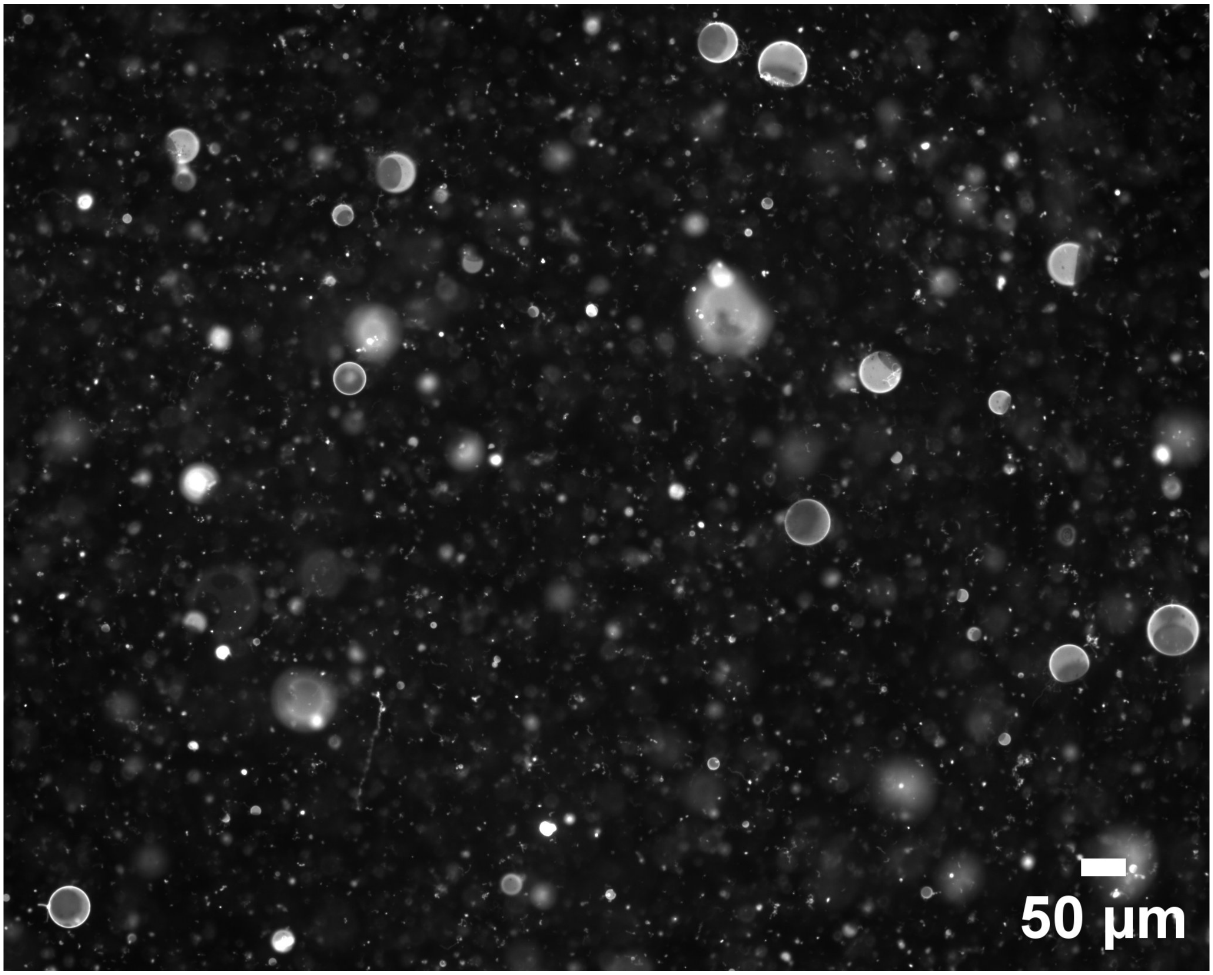

**Figure S2:** Representative wide field fluorescence micrograph of phase-separated GUVs at room temperature. The lipid composition corresponds to point 1 in Figure 2A of the main text. The vesicles were electroformed in 100 mM sucrose and diluted in an isotonic sucrose solution.

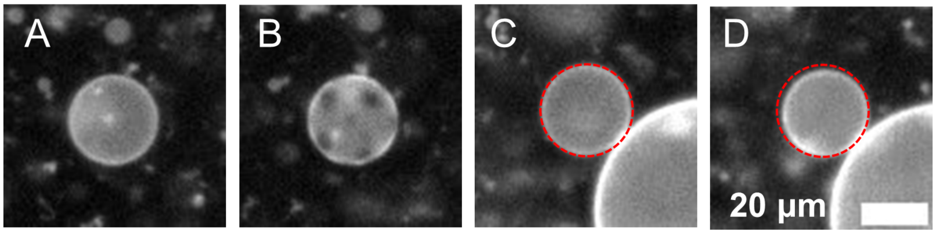

**Figure S3:** Fluorescence micrographs from a timelapse of a vesicle in a solution that was diluted immediately before the movie began. The vesicle membrane is initially uniform (A). Then the membrane phase separates into two liquid phases (B), becomes uniform again (C), and ruptures (D). Rupture is observed as an abrupt decrease in vesicle radius that occurs in one frame of the movie. Initially, all vesicles have tubes and aggregates (bright puncta in panels A and B). Fewer puncta are observed as rupture approaches. In the final frames of the movie, it is not clear if the vesicle is touching another vesicle, which might allow it to exchange lipids. However, if it were exchanging lipids, we might not expect to observe an abrupt decrease in radius that is characteristic of vesicle rupture. To highlight the size-change due to the rupture, the vesicle boundary is outlined with a red dotted line in panel C), and the same circle from Panel C is repeated in panel D). This figure corresponds to Supplemental Movie S3.

A)

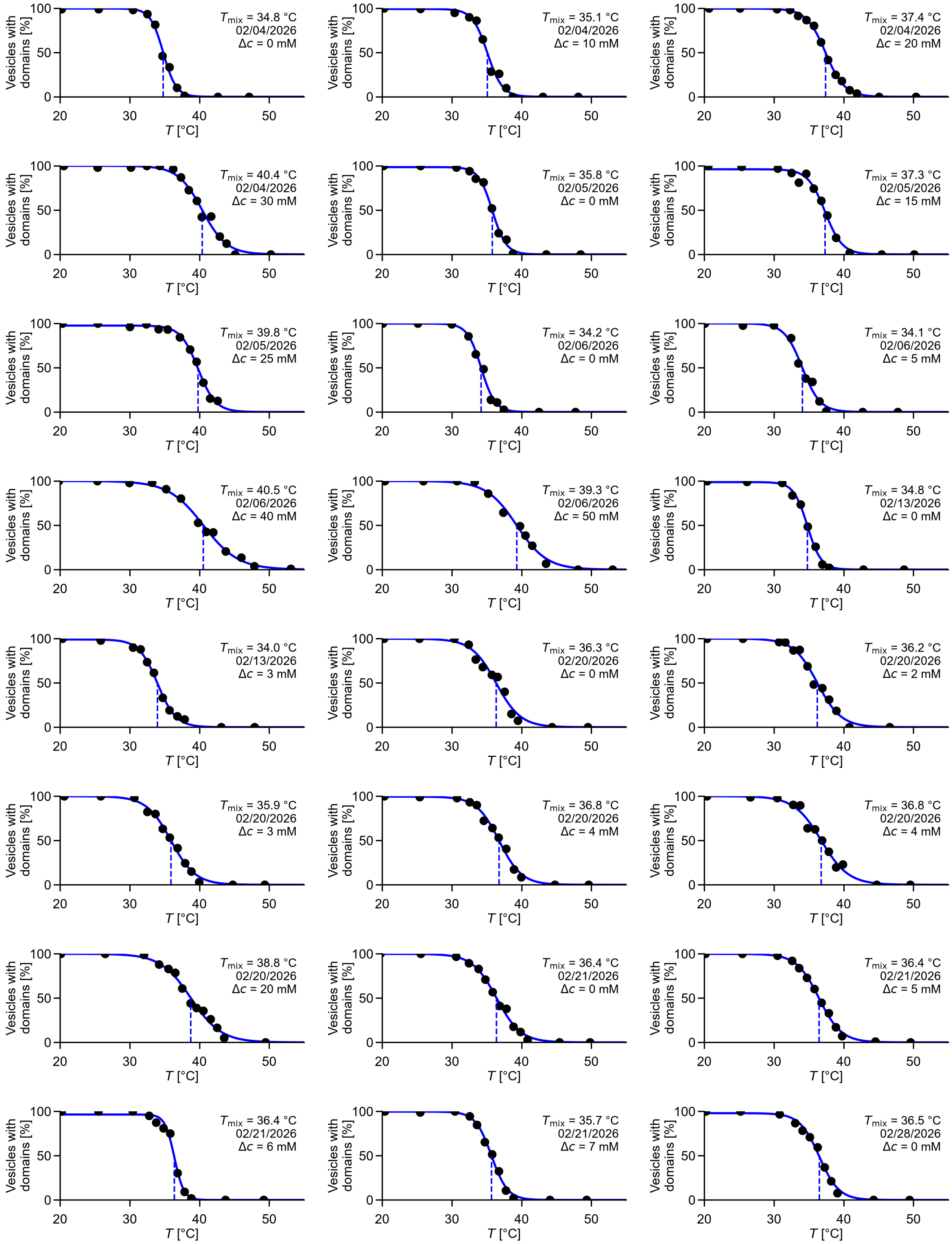
Vesicles with domains [%]
T [°C]
100
50
0
20
30
40
50
$T_{mix}$ = 34.8 °C
02/04/2026
Δc = 0 mM
$T_{mix}$ = 35.1 °C
02/04/2026
Δc = 10 mM
$T_{mix}$ = 37.4 °C
02/04/2026
Δc = 20 mM
$T_{mix}$ = 40.4 °C
02/04/2026
Δc = 30 mM
$T_{mix}$ = 35.8 °C
02/05/2026
Δc = 0 mM
$T_{mix}$ = 37.3 °C
02/05/2026
Δc = 15 mM
$T_{mix}$ = 39.8 °C
02/05/2026
Δc = 25 mM
$T_{mix}$ = 34.2 °C
02/06/2026
Δc = 0 mM
$T_{mix}$ = 34.1 °C
02/06/2026
Δc = 5 mM
$T_{mix}$ = 40.5 °C
02/06/2026
Δc = 40 mM
$T_{mix}$ = 39.3 °C
02/06/2026
Δc = 50 mM
$T_{mix}$ = 34.8 °C
02/13/2026
Δc = 0 mM
$T_{mix}$ = 34.0 °C
02/13/2026
Δc = 3 mM
$T_{mix}$ = 36.3 °C
02/20/2026
Δc = 0 mM
$T_{mix}$ = 36.2 °C
02/20/2026
Δc = 2 mM
$T_{mix}$ = 35.9 °C
02/20/2026
Δc = 3 mM
$T_{mix}$ = 36.8 °C
02/20/2026
Δc = 4 mM
$T_{mix}$ = 36.8 °C
02/20/2026
Δc = 4 mM
$T_{mix}$ = 38.8 °C
02/20/2026
Δc = 20 mM
$T_{mix}$ = 36.4 °C
02/21/2026
Δc = 0 mM
$T_{mix}$ = 36.4 °C
02/21/2026
Δc = 5 mM
$T_{mix}$ = 36.4 °C
02/21/2026
Δc = 6 mM
$T_{mix}$ = 35.7 °C
02/21/2026
Δc = 7 mM
$T_{mix}$ = 36.5 °C
02/28/2026
Δc = 0 mM

Continued...

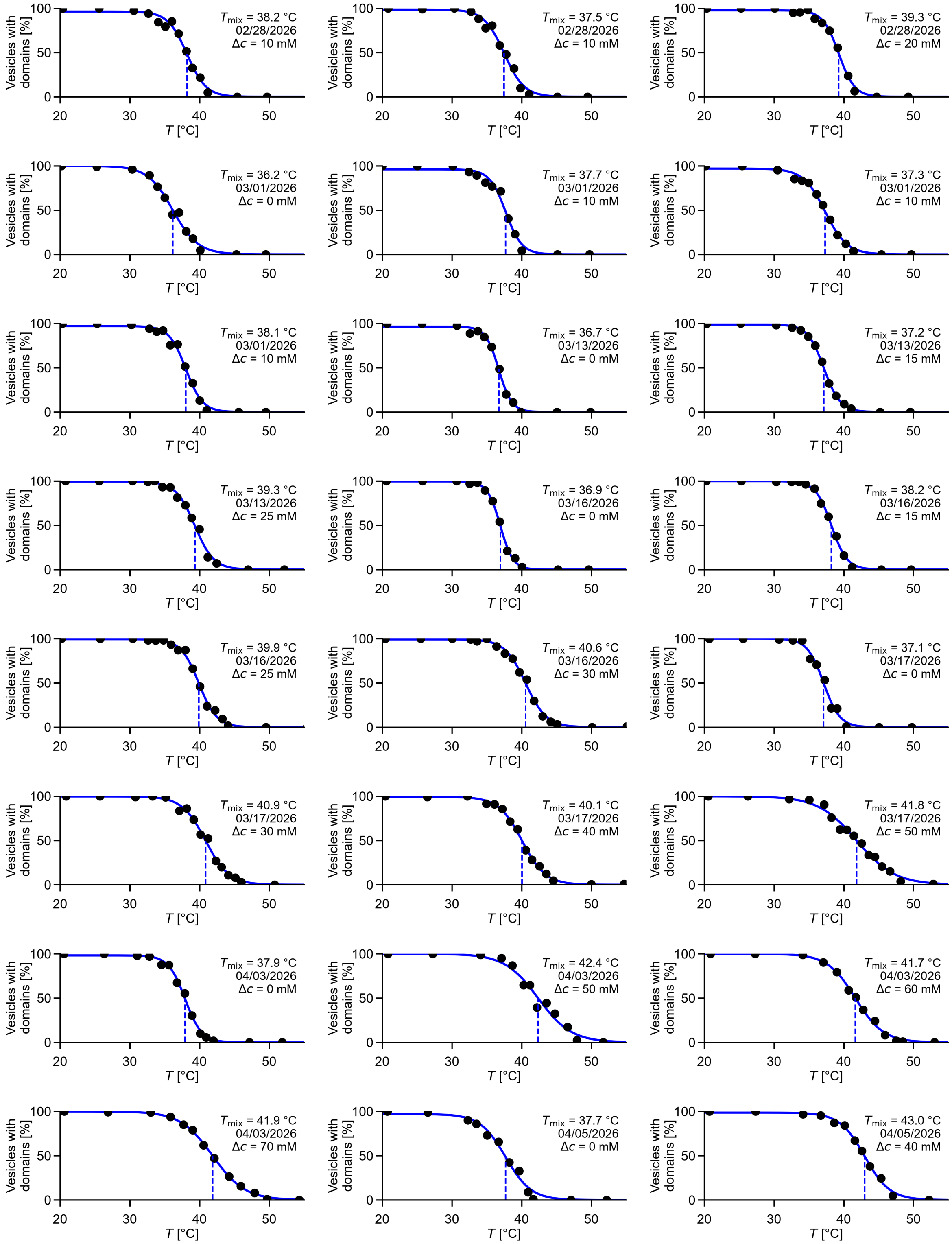
Vesicles with domains [%]
T [°C]
T_mix = 38.2 °C
02/28/2026
Δc = 10 mM
T_mix = 37.5 °C
02/28/2026
Δc = 10 mM
T_mix = 39.3 °C
02/28/2026
Δc = 20 mM
T_mix = 36.2 °C
03/01/2026
Δc = 0 mM
T_mix = 37.7 °C
03/01/2026
Δc = 10 mM
T_mix = 37.3 °C
03/01/2026
Δc = 10 mM
T_mix = 38.1 °C
03/01/2026
Δc = 10 mM
T_mix = 36.7 °C
03/13/2026
Δc = 0 mM
T_mix = 37.2 °C
03/13/2026
Δc = 15 mM
T_mix = 39.3 °C
03/13/2026
Δc = 25 mM
T_mix = 36.9 °C
03/16/2026
Δc = 0 mM
T_mix = 38.2 °C
03/16/2026
Δc = 15 mM
T_mix = 39.9 °C
03/16/2026
Δc = 25 mM
T_mix = 40.6 °C
03/16/2026
Δc = 30 mM
T_mix = 37.1 °C
03/17/2026
Δc = 0 mM
T_mix = 40.9 °C
03/17/2026
Δc = 30 mM
T_mix = 40.1 °C
03/17/2026
Δc = 40 mM
T_mix = 41.8 °C
03/17/2026
Δc = 50 mM
T_mix = 37.9 °C
04/03/2026
Δc = 0 mM
T_mix = 42.4 °C
04/03/2026
Δc = 50 mM
T_mix = 41.7 °C
04/03/2026
Δc = 60 mM
T_mix = 41.9 °C
04/03/2026
Δc = 70 mM
T_mix = 37.7 °C
04/05/2026
Δc = 0 mM
T_mix = 43.0 °C
04/05/2026
Δc = 40 mM

Continued...

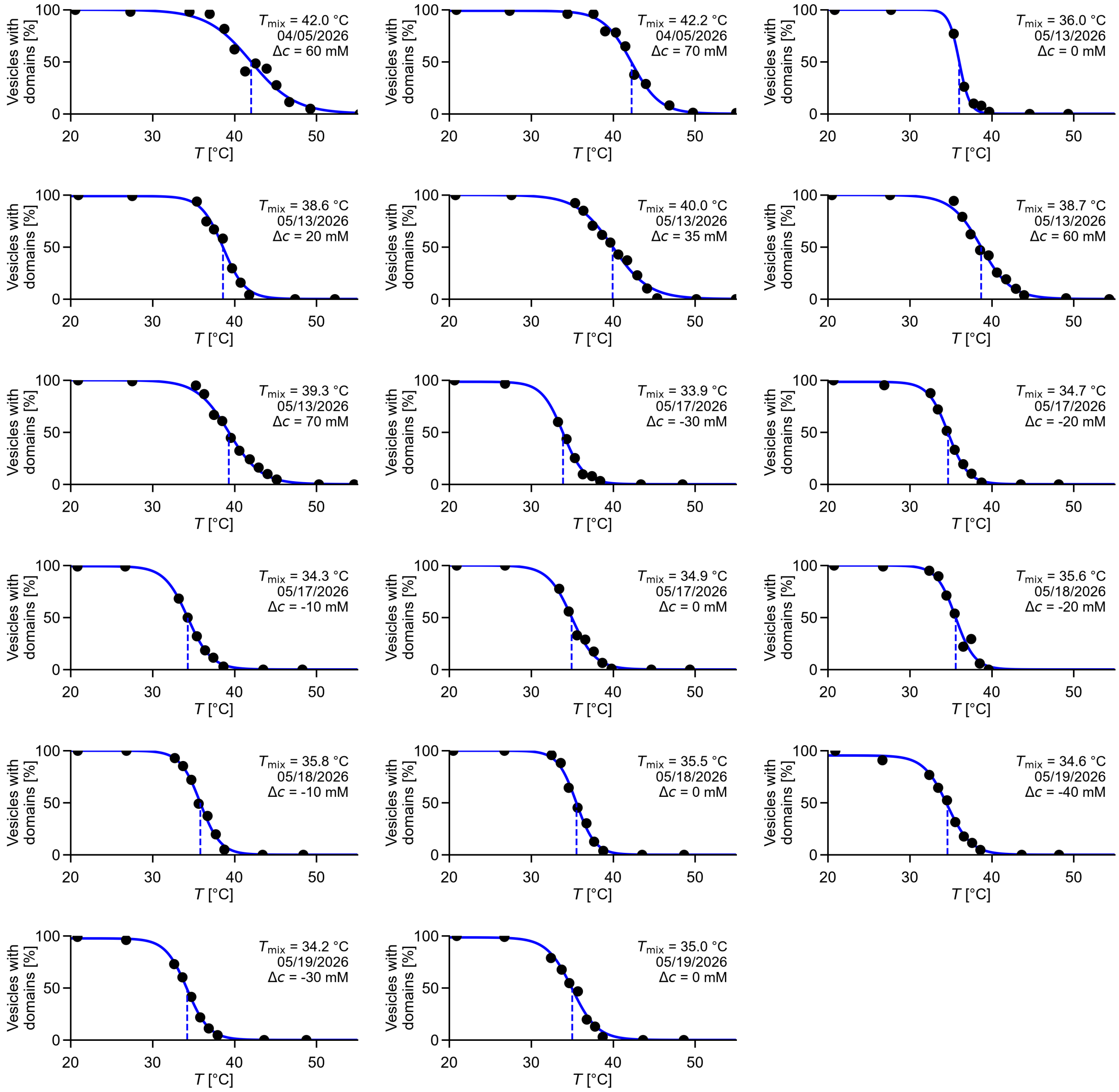

Continued...

B)

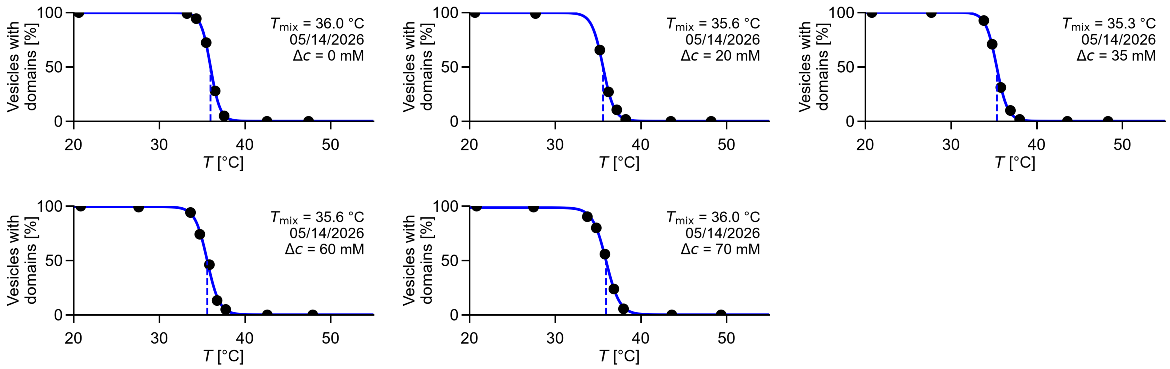

**Figure S4:** Data for the percentage of vesicles with coexisting liquid phases (as opposed to one uniform liquid phase) versus temperature at various osmolarity differences for a population of vesicles. Vesicles were electroformed in 100mM sucrose, using Lipid Ratio 1 in Figure 2 of the main text. A) Data were collected within 1 hour of imposing the osmolarity difference, and each transition temperature is plotted as a filled circle in Figure 4A of the main text. Each unique date (month/day/year) denotes a different electroformation batch of vesicles. B) Data were measured 21 hours after the dilution. Temperature values correspond to the open circles in Figure 4A of the main text. The vesicles used for this measurement were from the same electroformation batch as those used in the 05/13/2026 (month/day/year) measurement in panel A.

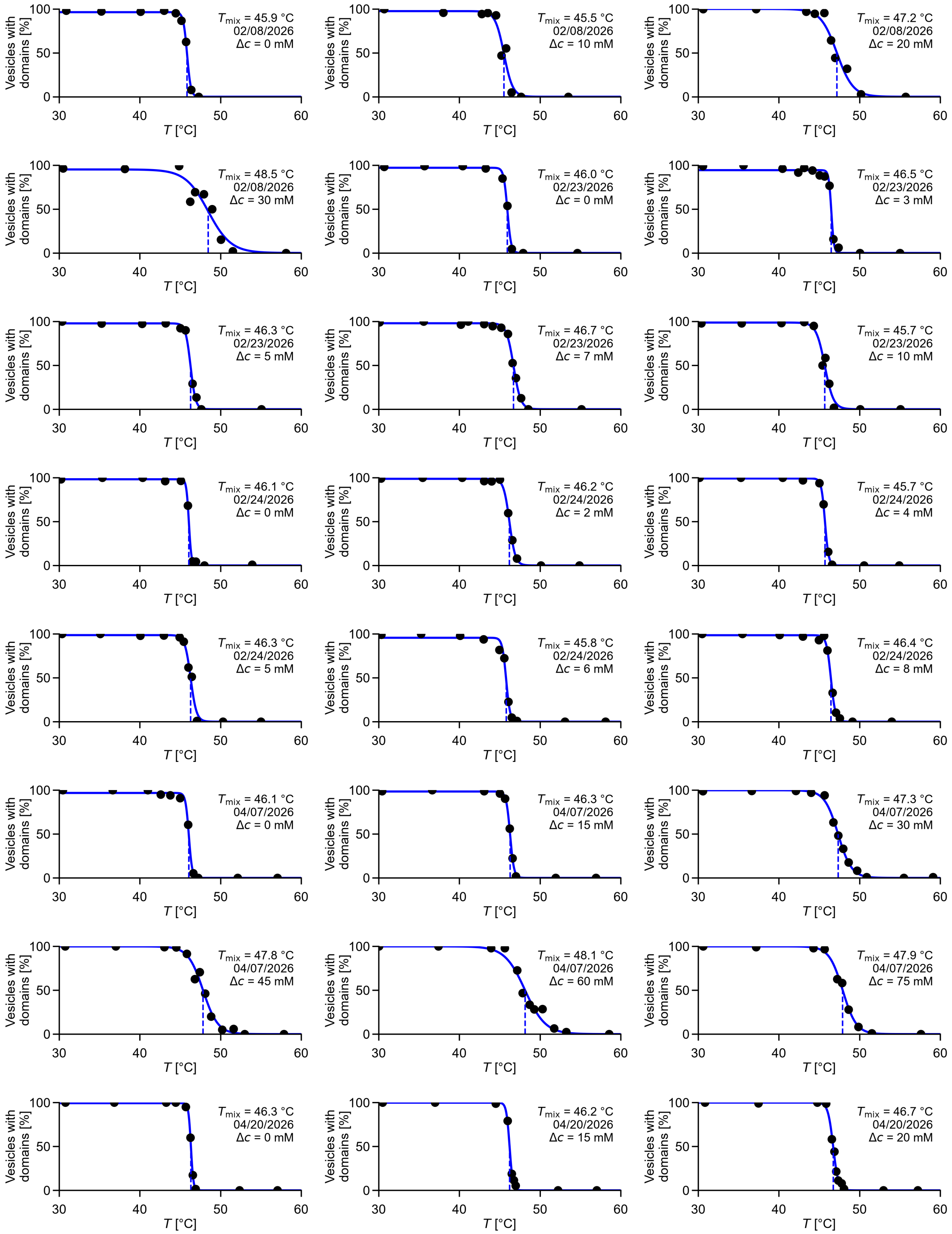
Vesicles with domains [%]
T [°C]
$T_{mix}$ = 45.9 °C 02/08/2026 Δc = 0 mM
$T_{mix}$ = 45.5 °C 02/08/2026 Δc = 10 mM
$T_{mix}$ = 47.2 °C 02/08/2026 Δc = 20 mM
$T_{mix}$ = 48.5 °C 02/08/2026 Δc = 30 mM
$T_{mix}$ = 46.0 °C 02/23/2026 Δc = 0 mM
$T_{mix}$ = 46.5 °C 02/23/2026 Δc = 3 mM
$T_{mix}$ = 46.3 °C 02/23/2026 Δc = 5 mM
$T_{mix}$ = 46.7 °C 02/23/2026 Δc = 7 mM
$T_{mix}$ = 45.7 °C 02/23/2026 Δc = 10 mM
$T_{mix}$ = 46.1 °C 02/24/2026 Δc = 0 mM
$T_{mix}$ = 46.2 °C 02/24/2026 Δc = 2 mM
$T_{mix}$ = 45.7 °C 02/24/2026 Δc = 4 mM
$T_{mix}$ = 46.3 °C 02/24/2026 Δc = 5 mM
$T_{mix}$ = 45.8 °C 02/24/2026 Δc = 6 mM
$T_{mix}$ = 46.4 °C 02/24/2026 Δc = 8 mM
$T_{mix}$ = 46.1 °C 04/07/2026 Δc = 0 mM
$T_{mix}$ = 46.3 °C 04/07/2026 Δc = 15 mM
$T_{mix}$ = 47.3 °C 04/07/2026 Δc = 30 mM
$T_{mix}$ = 47.8 °C 04/07/2026 Δc = 45 mM
$T_{mix}$ = 48.1 °C 04/07/2026 Δc = 60 mM
$T_{mix}$ = 47.9 °C 04/07/2026 Δc = 75 mM
$T_{mix}$ = 46.3 °C 04/20/2026 Δc = 0 mM
$T_{mix}$ = 46.2 °C 04/20/2026 Δc = 15 mM
$T_{mix}$ = 46.7 °C 04/20/2026 Δc = 20 mM

Continued...

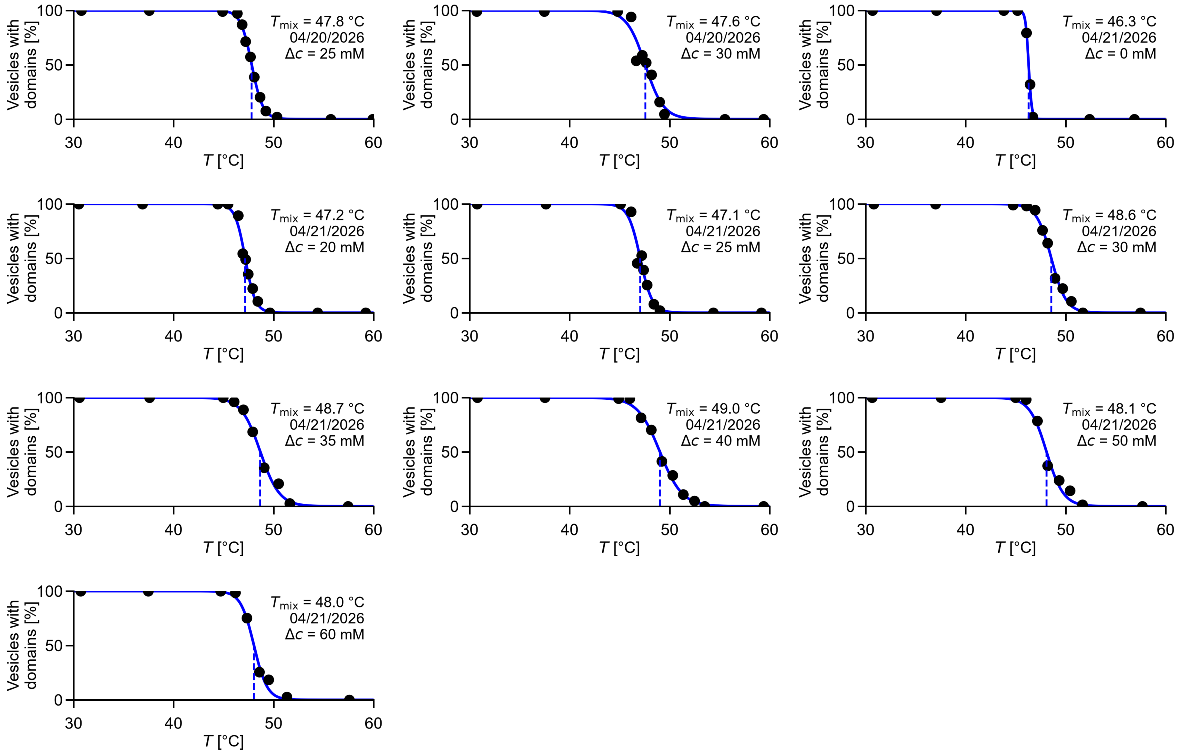

**Figure S5:** Data for the percentage of vesicles with coexisting liquid phases (as opposed to one uniform liquid phase) versus temperature at various osmolarity differences for a population of vesicles. Vesicles were electroformed in 100mM sucrose, using Lipid Ratio 2 in Figure 2 of the main text. Each unique date (month/day/year) denotes a different electroformation batch of vesicles. Data were collected within 1 hour of imposing the osmolarity difference, and each transition temperature is plotted as a circle in Figure 4B of the main text.

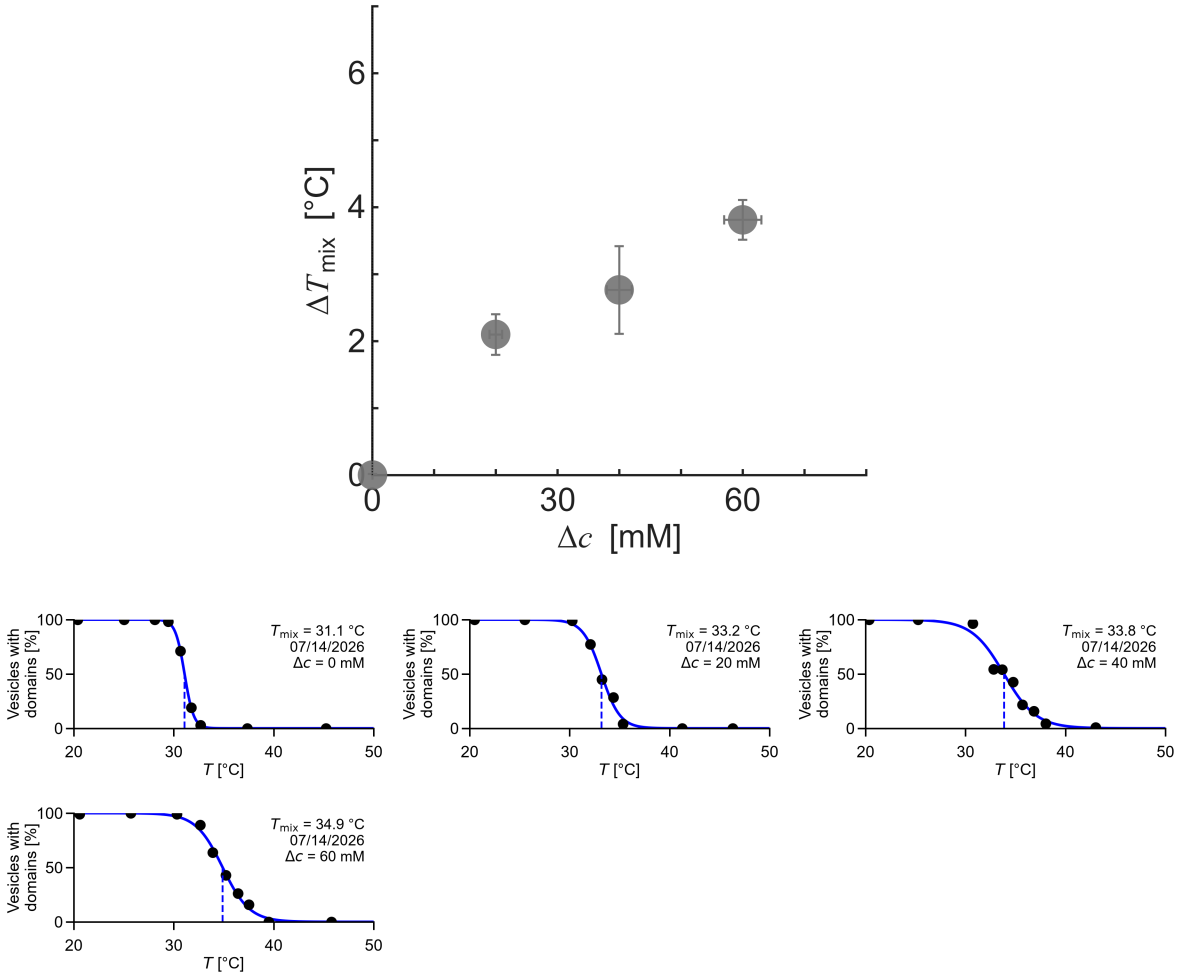

**Figure S6:** (Top) Increase in $T_{mix}$ versus osmolarity difference for a population of vesicles composed of 33/33/33 mol% DiPhyPC/DMPC/cholesterol, with 0.8 mol% dye (Rhodamine DPPE). Error bars are calculated as described in the Methods. (Bottom) Raw data for the four data points in the top panel. Each graph shows the percentage of vesicles with coexisting liquid phases (as opposed to one uniform liquid phase) versus temperature at various osmolarity differences for a population of vesicles. Vesicles were electroformed in 100 mM sucrose. Data were collected within 1 hour of imposing the osmolarity difference.

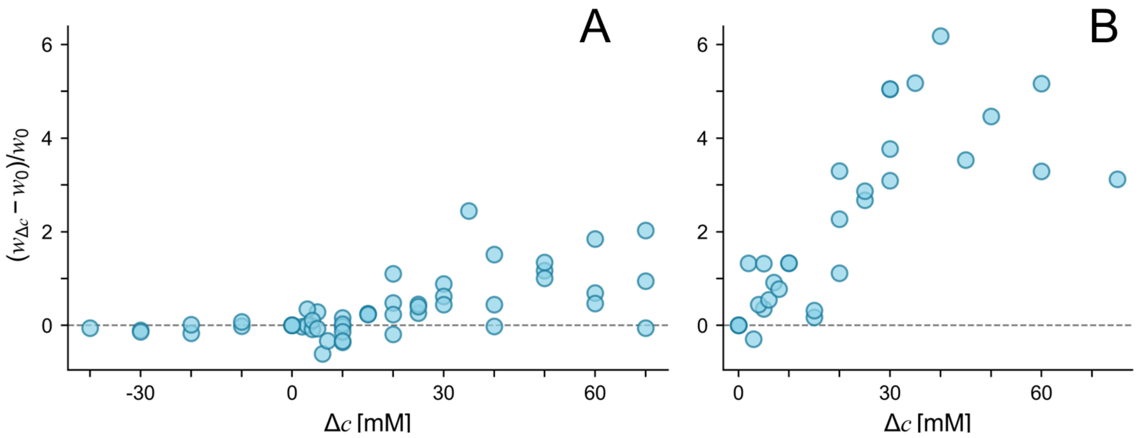

**Figure S7:** Relative widths of sigmoidal curves versus differences in osmolarity, corresponding to lipid compositions and data in Figure 4 of the main text. Relative widths are defined as $(w_{\Delta c} - w_0)/w_0$. Widths, $w$, are defined by the equation *Percent phase-separated* = *Max* $[1 - (1 + \exp(-(T - T_{\text{mix}})/w))^{-1}]$. $w_{\Delta c}$ is the width at $\Delta c \neq 0$, and $w_0$ is the width at $\Delta c = 0$ from the same sample. In other words, each relative width was calculated from a pair of values ($w_{\Delta c}$ and $w_0$) measured from vesicles produced in the same electroformation sample, to account for day-to-day variation. In both panels, relative widths increase with positive values of $\Delta c$ and reach maximums before the plateaus in Figure 4 of the main text. Larger relative widths in Panel B arise because $w_0$ in composition 2 is small.

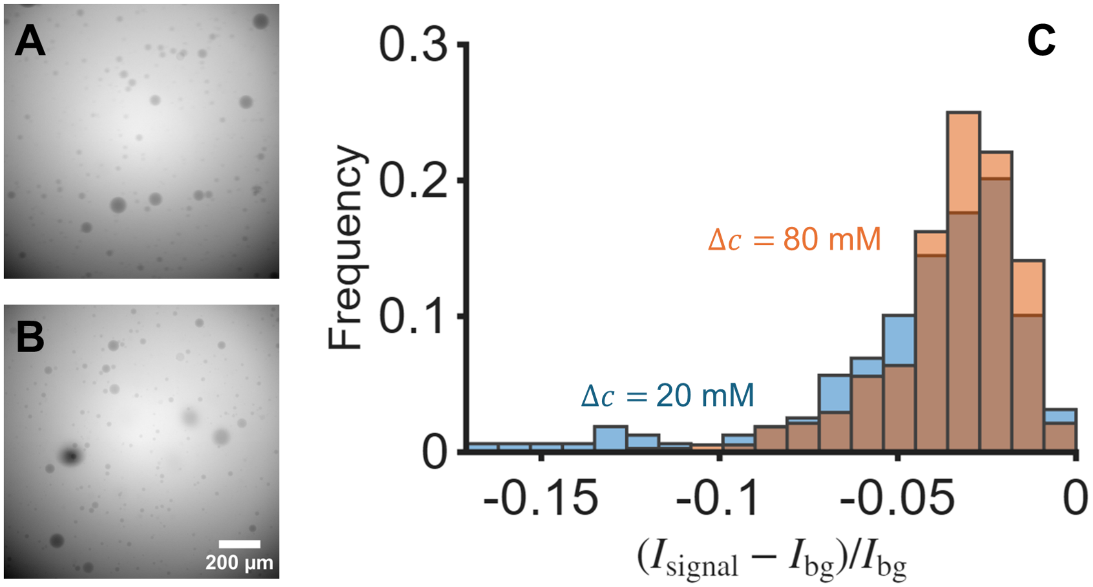

**Figure S8:** Evaluation of the use of a calcein-based assay to determine whether giant unilamellar vesicles have ruptured. The lipid ratio corresponds to point 1 in Figure 2A of the main text. Vesicles were electroformed in 100 mM sucrose and diluted with a lower-concentration sucrose solution to impose concentration difference of either 20 mM or 80 mM. Also, a calcein aqueous solution was added to a final calcein concentration of 50 µM. The dilution caused by the aqueous calcein solution was included when calculating the final sucrose concentrations. Fluorescence images of calcein were taken at 25 °C under each osmotic condition. Panel A) and B) show representative images from concentration differences of 20 mM and 80 mM, respectively. For each vesicle, the apparent fluorescence intensity of the dark vesicle $I_{signal}$ and that of the immediately adjacent background $I_{bg}$ were measured as a pair. A total of 159 and 376 intensity pairs were collected for $\Delta c$ = 20 mM and 80 mM, respectively. For each pair, the relative brightness of the vesicle compared with the surrounding background $(I_{signal}–I_{bg})/I_{bg}$ was calculated. (C) Although very dark vesicle regions were less frequent at $\Delta c$ = 80 mM than at $\Delta c$ = 20 mM, implying that calcein had entered those vesicles, the overall distribution changed little.

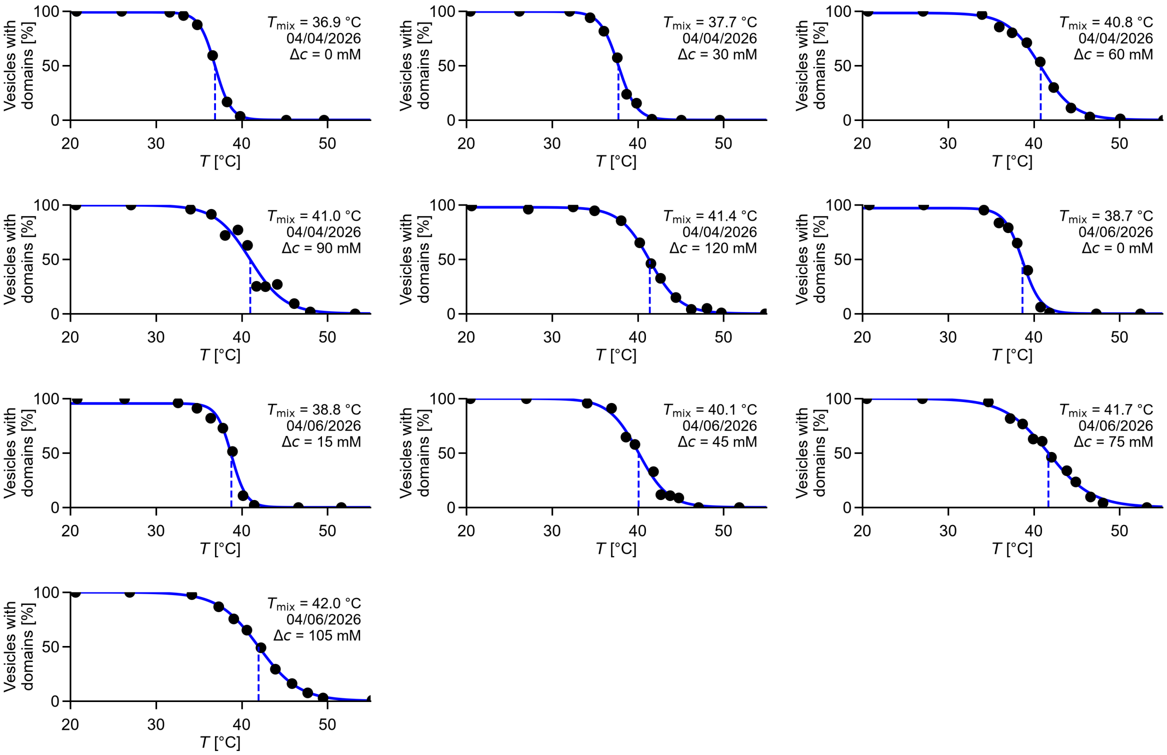

**Figure S9:** Data for the percentage of vesicles with coexisting liquid phases (as opposed to one uniform liquid phase) versus temperature at various osmolarity differences for a population of vesicles. Vesicles were electroformed in 150 mM sucrose, using Lipid Ratio 1 in Figure 2A of the main text. Data were collected within 1 hour of imposing the osmolarity difference, and each transition temperature is plotted as a triangle in Figure 6A of the main text.

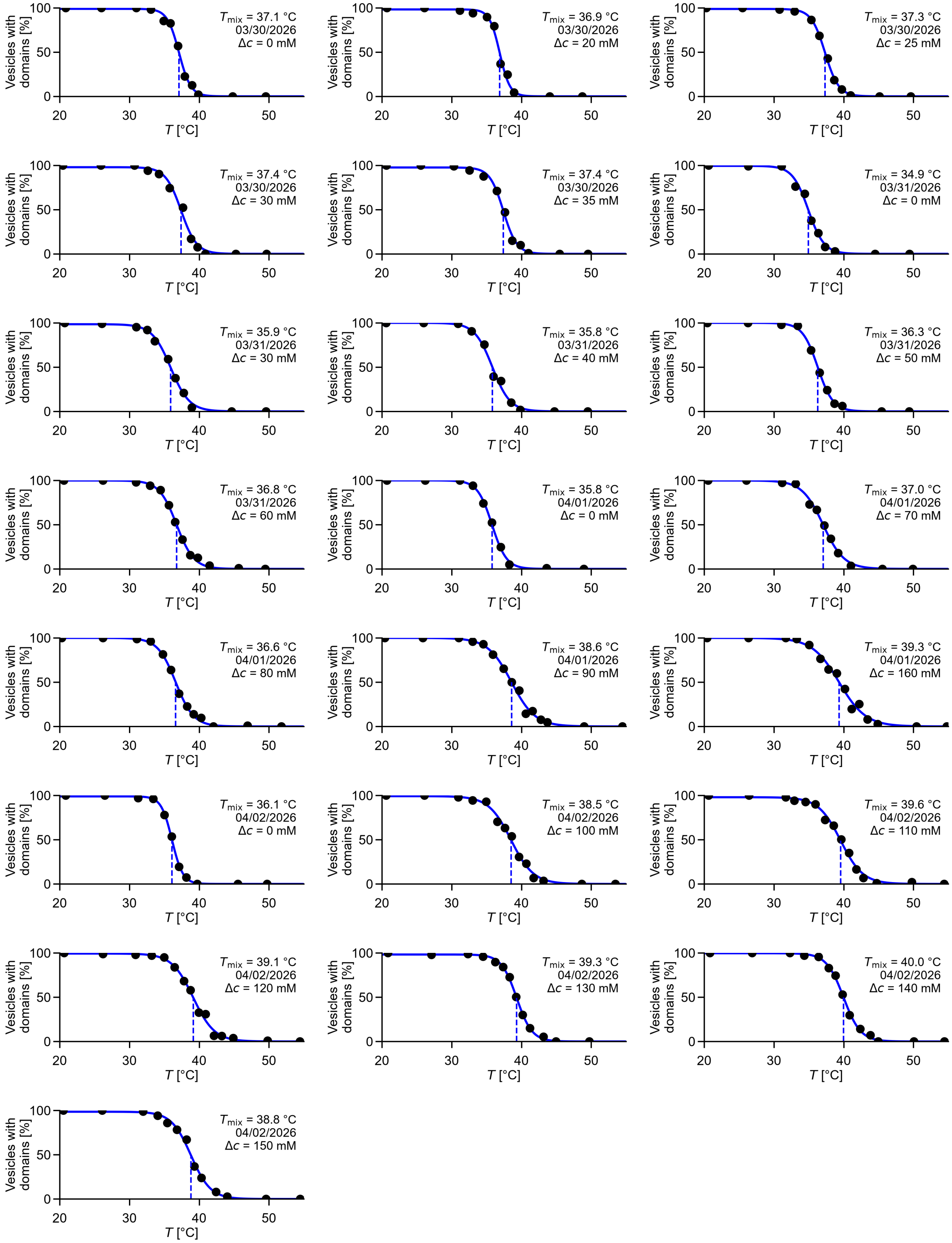
Vesicles with domains [%]
T [°C]
$T_{mix}$ = 37.1 °C 03/30/2026 Δc = 0 mM
$T_{mix}$ = 36.9 °C 03/30/2026 Δc = 20 mM
$T_{mix}$ = 37.3 °C 03/30/2026 Δc = 25 mM
$T_{mix}$ = 37.4 °C 03/30/2026 Δc = 30 mM
$T_{mix}$ = 37.4 °C 03/30/2026 Δc = 35 mM
$T_{mix}$ = 34.9 °C 03/31/2026 Δc = 0 mM
$T_{mix}$ = 35.9 °C 03/31/2026 Δc = 30 mM
$T_{mix}$ = 35.8 °C 03/31/2026 Δc = 40 mM
$T_{mix}$ = 36.3 °C 03/31/2026 Δc = 50 mM
$T_{mix}$ = 36.8 °C 03/31/2026 Δc = 60 mM
$T_{mix}$ = 35.8 °C 04/01/2026 Δc = 0 mM
$T_{mix}$ = 37.0 °C 04/01/2026 Δc = 70 mM
$T_{mix}$ = 36.6 °C 04/01/2026 Δc = 80 mM
$T_{mix}$ = 38.6 °C 04/01/2026 Δc = 90 mM
$T_{mix}$ = 39.3 °C 04/01/2026 Δc = 160 mM
$T_{mix}$ = 36.1 °C 04/02/2026 Δc = 0 mM
$T_{mix}$ = 38.5 °C 04/02/2026 Δc = 100 mM
$T_{mix}$ = 39.6 °C 04/02/2026 Δc = 110 mM
$T_{mix}$ = 39.1 °C 04/02/2026 Δc = 120 mM
$T_{mix}$ = 39.3 °C 04/02/2026 Δc = 130 mM
$T_{mix}$ = 40.0 °C 04/02/2026 Δc = 140 mM
$T_{mix}$ = 38.8 °C 04/02/2026 Δc = 150 mM

Continued...

**Figure S10 (Continued from previous page):** Data for the percentage of vesicles with coexisting liquid phases (as opposed to one uniform liquid phase) versus temperature at various osmolarity differences for a population of vesicles. Vesicles were electroformed in 200 mM sucrose, using Lipid Ratio 1 in Figure 2A of the main text. Data were collected within 1 hour of imposing the osmolarity difference, and each transition temperature is plotted as a diamond in Figure 6A of the main text.

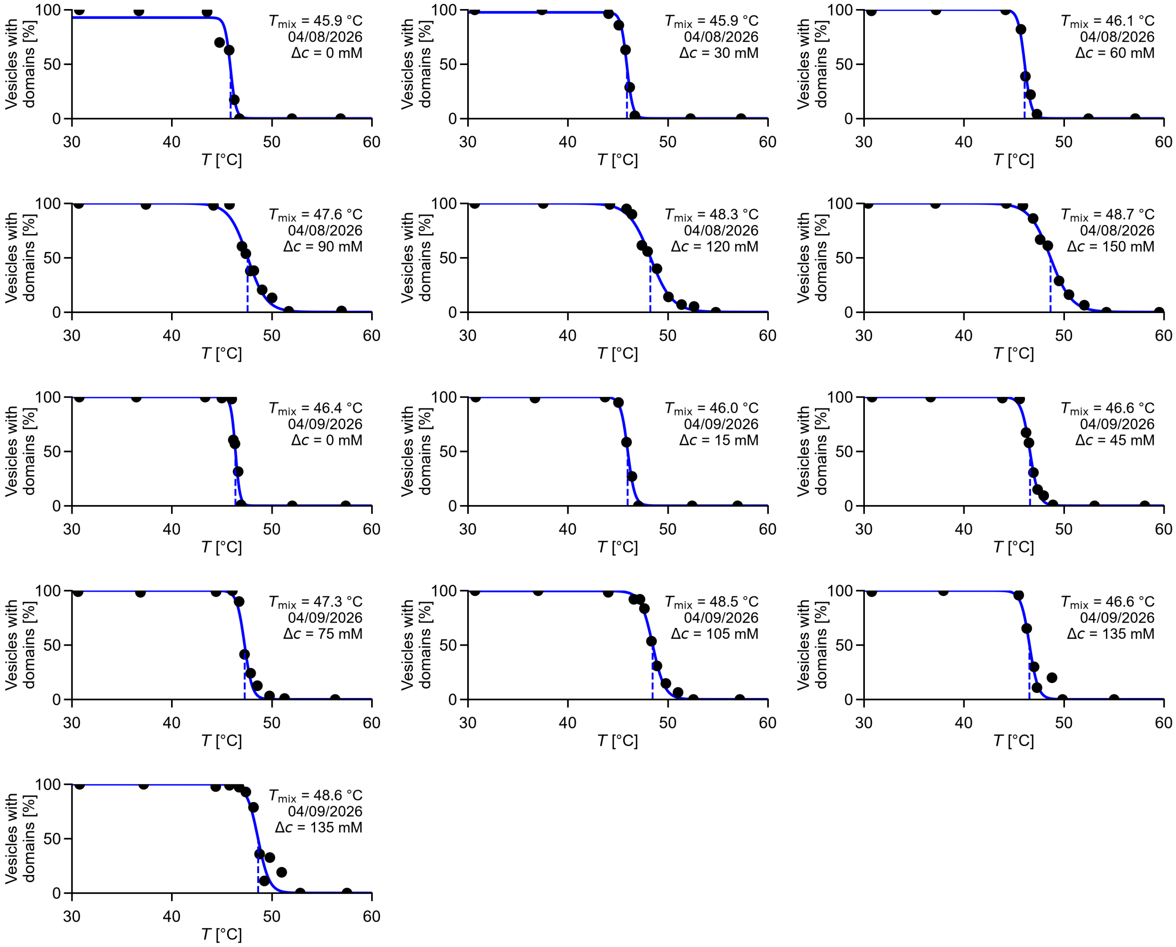

**Figure S11:** Data for the percentage of vesicles with coexisting liquid phases (as opposed to one uniform liquid phase) versus temperature at various osmolarity differences for a population of vesicles. Vesicles were electroformed in 200 mM sucrose, using Lipid Ratio 2 in Figure 2A of the main text. Data were collected within 1 hour of imposing the osmolarity difference, and each transition temperature is plotted as a diamond in Figure 6B of the main text.

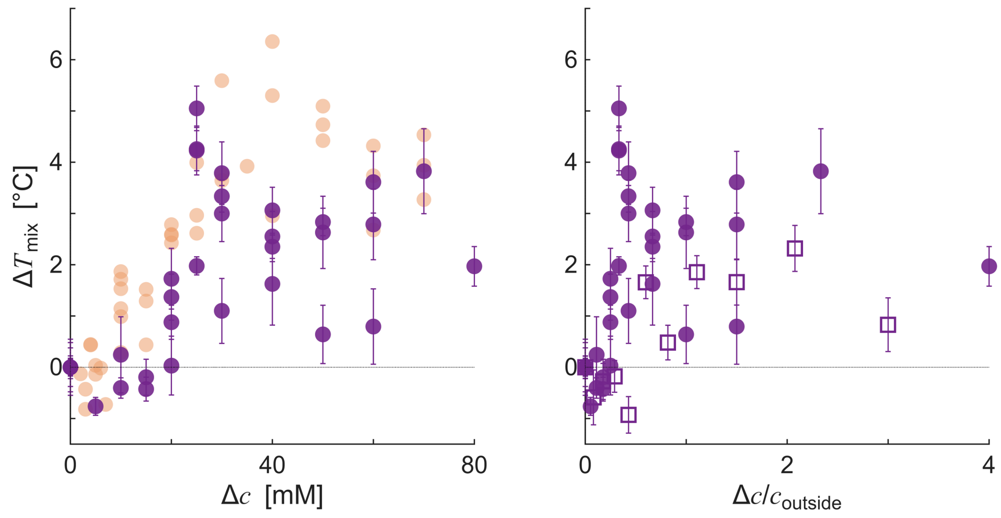

**Figure S12:** Shift in liquid-liquid phase transition temperatures ($\Delta T_{mix}$) for a population of GUV membranes versus the osmolarity difference ($\Delta c$) with TMAO as an osmolyte instead of sucrose. The lipid ratio corresponds to point 1 in Figure 2A of the main text. (Left) Data for vesicles electroformed in 100 mM TMAO and diluted with a lower concentration of TMAO solution (dark purple filled circles). Data were taken within 1 hour of applying the osmolarity difference. For comparison, the 100 mM sucrose data from Figure 4A are overlaid as light orange filled circles. (Right) Rescaled plot of 100 mM TMAO data (dark purple filled circles) and 200 mM TMAO data (purple open squares) with the x-axis re-plotted as the relative concentration difference, $\Delta c/c_{outside}$. Vertical error bars are calculated as described in the Methods.

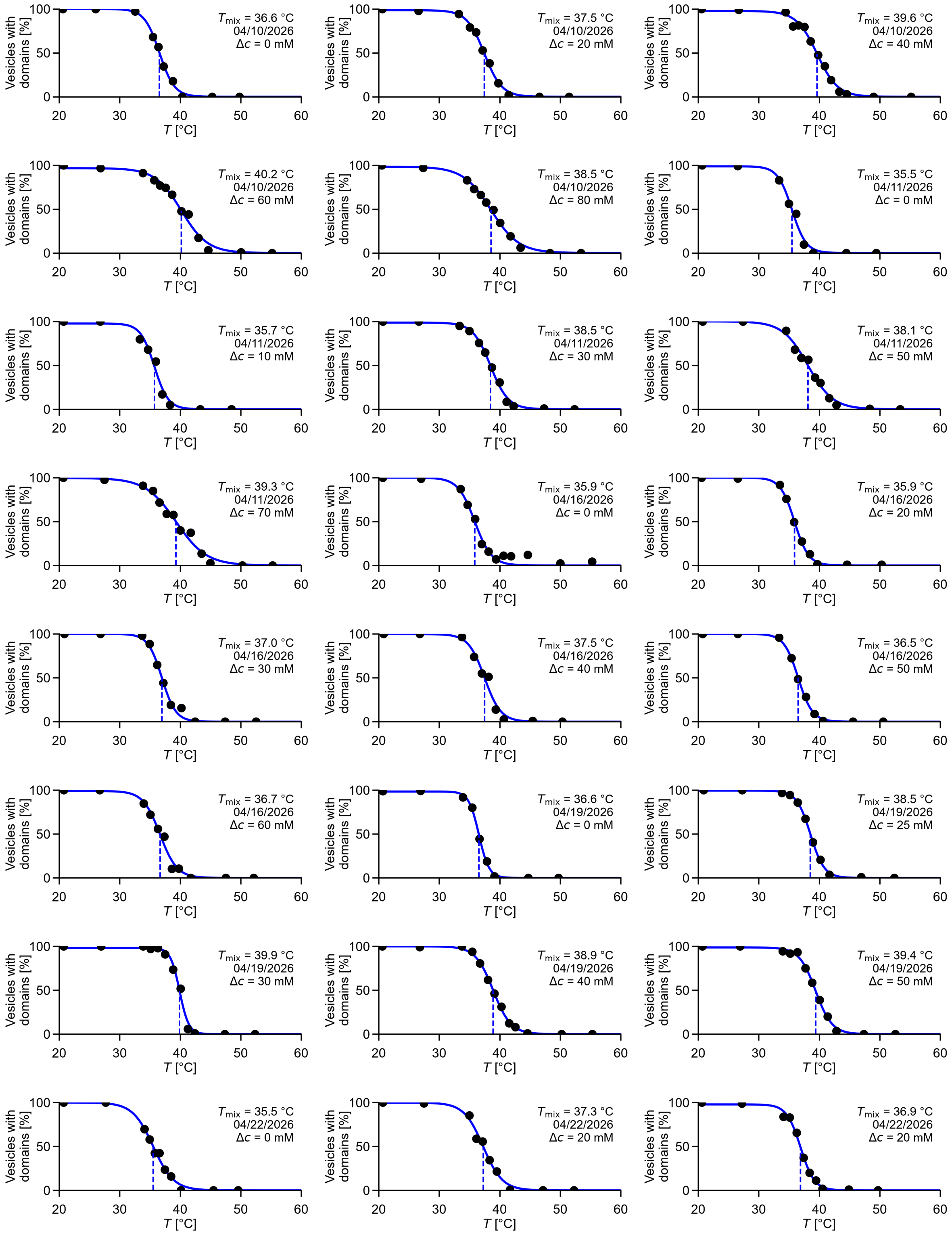
$T_{mix}$ = 36.6 °C
04/10/2026
$\Delta c$ = 0 mM
$T_{mix}$ = 37.5 °C
04/10/2026
$\Delta c$ = 20 mM
$T_{mix}$ = 39.6 °C
04/10/2026
$\Delta c$ = 40 mM
$T_{mix}$ = 40.2 °C
04/10/2026
$\Delta c$ = 60 mM
$T_{mix}$ = 38.5 °C
04/10/2026
$\Delta c$ = 80 mM
$T_{mix}$ = 35.5 °C
04/11/2026
$\Delta c$ = 0 mM
$T_{mix}$ = 35.7 °C
04/11/2026
$\Delta c$ = 10 mM
$T_{mix}$ = 38.5 °C
04/11/2026
$\Delta c$ = 30 mM
$T_{mix}$ = 38.1 °C
04/11/2026
$\Delta c$ = 50 mM
$T_{mix}$ = 39.3 °C
04/11/2026
$\Delta c$ = 70 mM
$T_{mix}$ = 35.9 °C
04/16/2026
$\Delta c$ = 0 mM
$T_{mix}$ = 35.9 °C
04/16/2026
$\Delta c$ = 20 mM
$T_{mix}$ = 37.0 °C
04/16/2026
$\Delta c$ = 30 mM
$T_{mix}$ = 37.5 °C
04/16/2026
$\Delta c$ = 40 mM
$T_{mix}$ = 36.5 °C
04/16/2026
$\Delta c$ = 50 mM
$T_{mix}$ = 36.7 °C
04/16/2026
$\Delta c$ = 60 mM
$T_{mix}$ = 36.6 °C
04/19/2026
$\Delta c$ = 0 mM
$T_{mix}$ = 38.5 °C
04/19/2026
$\Delta c$ = 25 mM
$T_{mix}$ = 39.9 °C
04/19/2026
$\Delta c$ = 30 mM
$T_{mix}$ = 38.9 °C
04/19/2026
$\Delta c$ = 40 mM
$T_{mix}$ = 39.4 °C
04/19/2026
$\Delta c$ = 50 mM
$T_{mix}$ = 35.5 °C
04/22/2026
$\Delta c$ = 0 mM
$T_{mix}$ = 37.3 °C
04/22/2026
$\Delta c$ = 20 mM
$T_{mix}$ = 36.9 °C
04/22/2026
$\Delta c$ = 20 mM
Vesicles with domains [%]
0
50
100
$T$ [°C]
20
30
40
50
60

Continued...

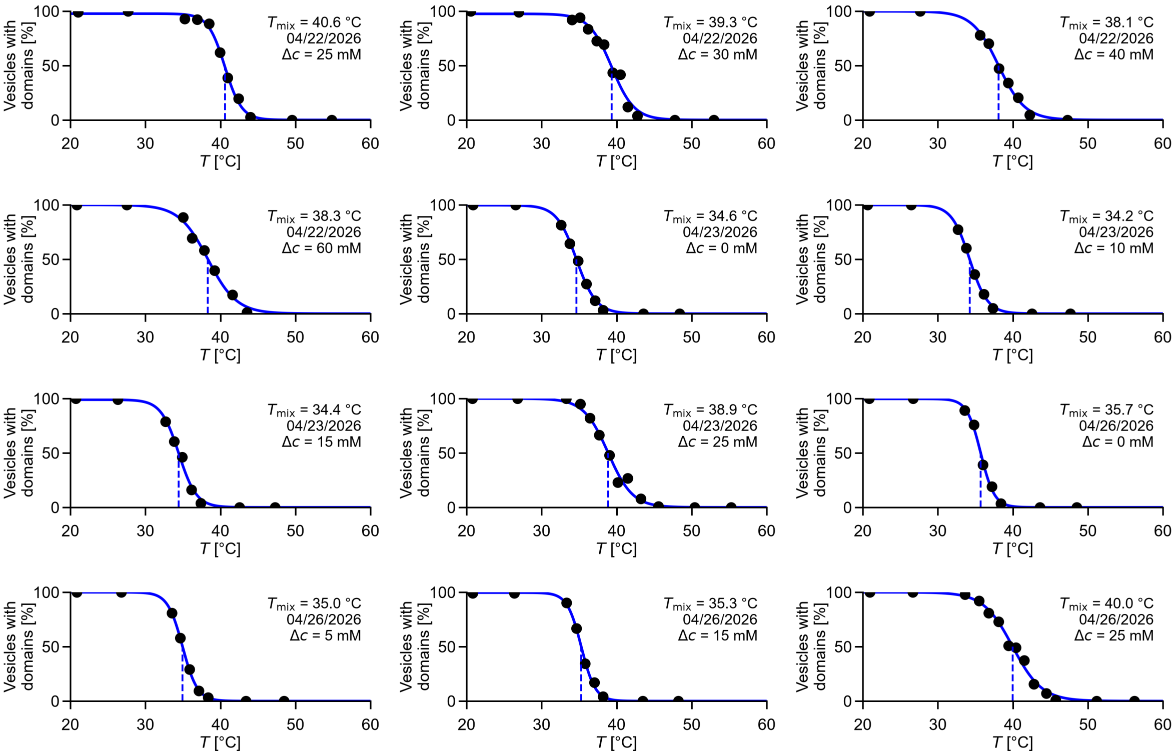

**Figure S13:** Data for the percentage of vesicles with coexisting liquid phases (as opposed to one uniform liquid phase) versus temperature at various osmolarity differences for a population of vesicles. Vesicles were electroformed in 100 mM TMAO, using Lipid Ratio 1 in Figure 2A of the main text. Data were collected within 1 hour of imposing the osmolarity difference, and each transition temperature is plotted as a purple filled circle in Supplemental Figure S12.

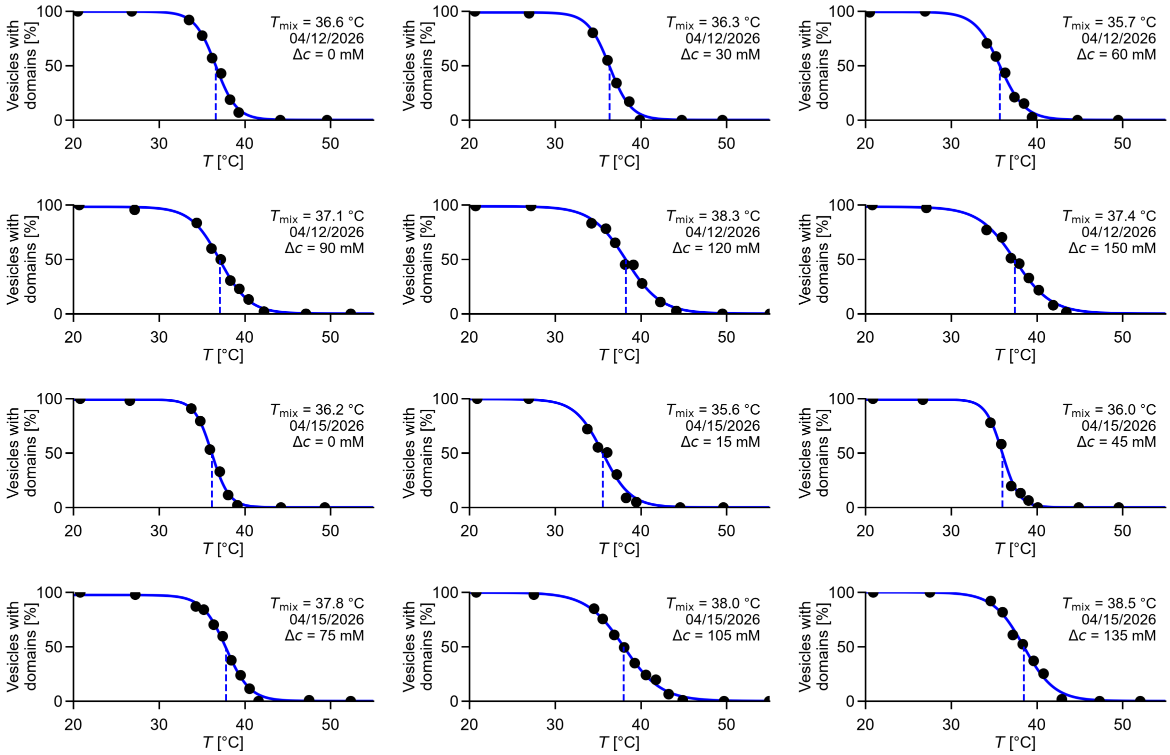

**Figure S14:** Data for the percentage of vesicles with coexisting liquid phases (as opposed to one uniform liquid phase) versus temperature at various osmolarity differences for a population of vesicles. Vesicles were electroformed in 200 mM TMAO, using Lipid Ratio 1 in Figure 2A of the main text. Data were collected within 1 hour of imposing the osmolarity difference, and each transition temperature is plotted as a purple open square in Supplemental Figure S12.

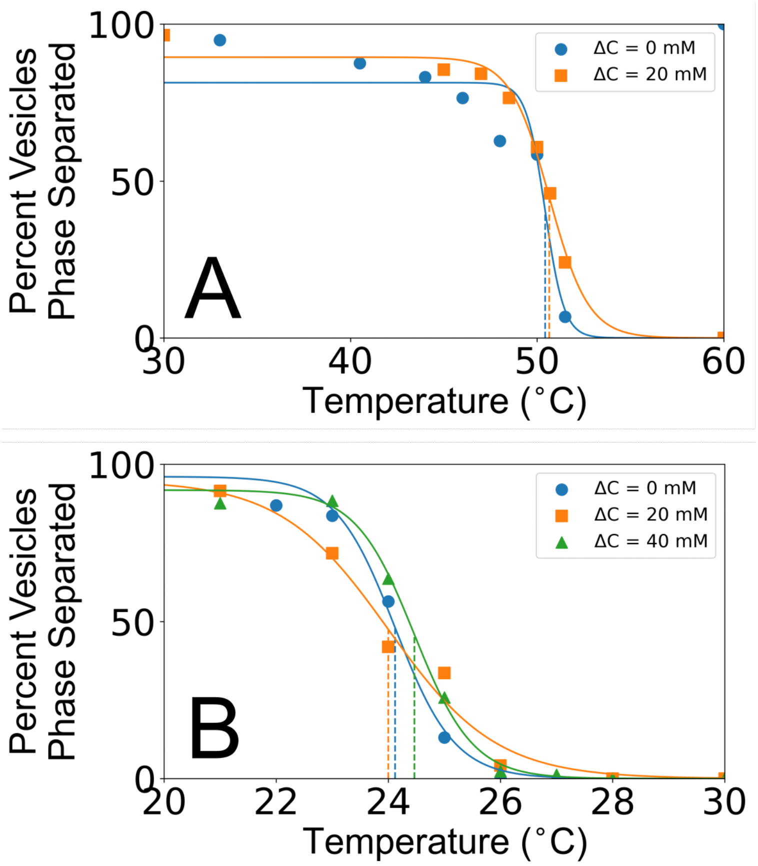

**Figure S15:** Application of osmotic pressure does not decrease $T_{\text{mix}}$ in vesicles of two additional lipid compositions. (A) Percent of vesicle membranes with coexisting liquid phases (instead of one uniform liquid phase) versus temperature for a population of vesicles composed of 6/44/40/10 mol% DiPhyPC/DPPC/cholesterol/DPPS, with 0.8 mol% dye (Rhodamine DPPE), at two osmolarity differences. Vesicles were electroformed in 100 mM sucrose. (B) Percent of vesicle membranes with coexisting liquid phases (instead of one uniform liquid phase) versus temperature for a population of vesicles composed of 6/54/40 mol% DiPhyPC/Di(13:0)PC/cholesterol, with 0.8 mol% dye (Rhodamine DPPE), at three osmolarity differences. Vesicles were electroformed in 200 mM sucrose.

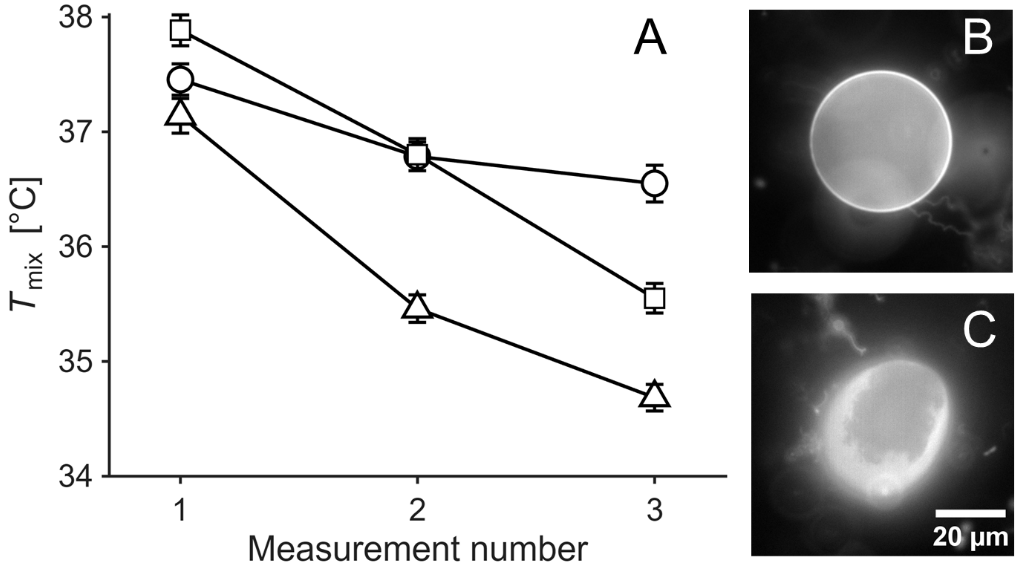

**Figure S16:** Liquid-liquid phase transition temperatures decrease with repeated heating and cooling cycles. The lipid ratio corresponds to point 1 in Figure 2A of the main text. (A) Transition temperatures of three vesicles (each shown by a different symbol), for three consecutive measurements. Transition temperatures were determined as vesicles were heated. Transition temperatures were defined as the mean of the highest temperature at which liquid domains were visible and the lowest temperature at which no domains were visible, as the vesicle was heated. Error bars represent half of the interval between these two temperatures. For each measurement, temperature sweeps were implemented in two consecutive stages. First, the temperature was raised from 20 °C to around 35 °C in one step. Next, the temperature was raised in increments of ~0.3 °C, with equilibration at 1 min at each temperature. After domains disappeared, the temperature was immediately returned to 20 °C. The sample was then allowed to equilibrate for 30 min before the second measurement, allowing domains to coalesce. The same procedure was followed for the third measurement. Lines in the plot connect points and are not fits. (B, C) Some vesicles became flaccid and distorted after repeated cooling–heating cycles. Panel B shows an image of one vesicle taken at 36 °C during the first measurement, and Panel C shows the same vesicle at 35 °C during the second measurement. In Panel C, the edges of the domain are fluctuating in a manner characteristic of miscibility critical points.

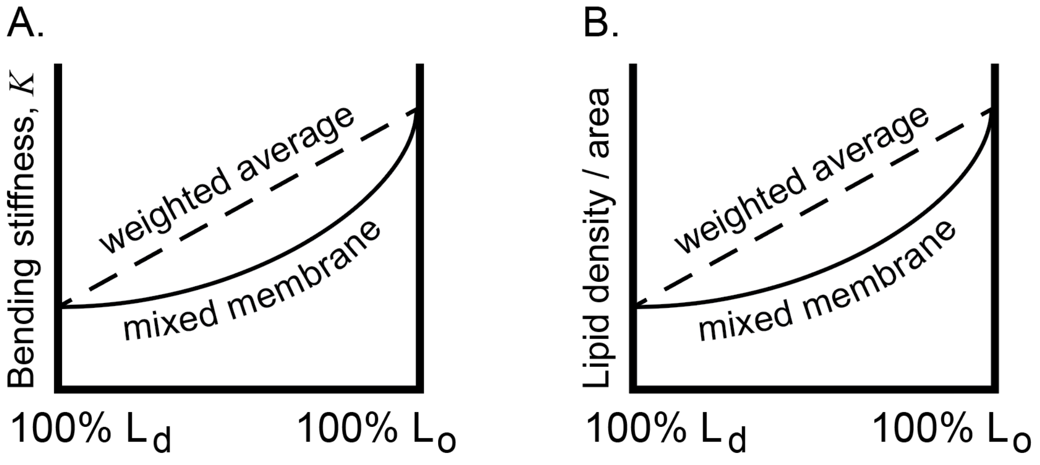

**Figure S17:** Schematic illustrating how the bending stiffness of a membrane (A) and its lipid density per unit area (B) are higher in a membrane of 100% Lo phase (right axis) than the in a membrane of 100% Ld phase (left axis). The values for a uniformly mixed, liquid membrane are lower than for a weighted average of the separate phases.

| Vesicle number | Radius at t = 0 [μm] | Radius at t = 5 min [μm] | Δradius [%] |
|---|---|---|---|
| 1 | 17.6 | 19.6 | 11.2 |
| 2 | 21.4 | 23.7 | 11.0 |
| 3 | 30.4 | 33.1 | 8.7 |
| 4 | 23.8 | 26.6 | 11.5 |
| 5 | 25.8 | 27.2 | 5.6 |
| 6 | 25.9 | 28.7 | 10.6 |

**Supplemental Table S1:** The change in radius of giant unilamellar vesicles due to dilution of the external solution. The radius of each vesicle was measured immediately before the dilution, corresponding to $t = 0$ in Supplemental Movie S2, and 5 min later. Dilution began approximately 7 s after the start of video acquisition. Experimental details and the corresponding video are provided in Supplemental Movie S2.

| No. | Vesicle radius [μm] | Micropipette radius [μm] | Suction pressure [kPa] | Transition tension [mN/m] | Temp [°C] | Ratio |
|---|---|---|---|---|---|---|
| 1 | 10.59 ± 0.14 | 3.03 ± 0.14 | 0.88 ± 0.12 | 1.9 ± 0.3 | 28 | 3 |
| 2 | 19.25 ± 0.14 | 3.71 ± 0.14 | 2.95 ± 0.58 | 6.8 ± 1.4 | 20 | 4 |
| 3 | 33.69 ± 0.14 | 3.71 ± 0.14 | 3.26 ± 0.61 | 6.8 ± 1.3 | 20 | 4 |
| 4 | 13.62 ± 0.14 | 3.71 ± 0.14 | 0.55 ± 0.30 | 1.4 ± 0.8 | 25 | 4 |
| 5 | 8.53 ± 0.14 | 3.71 ± 0.14 | 2.00 ± 0.25 | 6.6 ± 0.9 | 27 | 4 |
| 6 | 9.77 ± 0.14 | 3.71 ± 0.14 | 0.73 ± 0.09 | 2.2 ± 0.3 | 27 | 4 |

**Supplemental Table S2:** Experimental values and calculated tensions at the Lo-Ld transition of the aspirated vesicles in Figure 3F of the main text. Supplemental Movies S4 and S5 show vesicles #1 and #2, respectively, during aspiration. Tension was calculated as described in Methods. Uncertainties in the vesicle and micropipette radii represent the pixel resolution of images. The uncertainty in determining the onset of phase separation manifests as an uncertainty in the suction pressure. All uncertainties were propagated to determine the uncertainty in transition tension. Vesicle 1 has a lipid composition corresponding to point 3 in Figure 2A. Vesicles 2-6 were from the same electroformation batch with a lipid composition corresponding to point 4 in Figure 2A of the main text. Vesicles were electroformed in 100 mM sucrose and sank to the bottom of a chamber containing a 1:2 mixture of 100 mM sucrose and 100 mM glucose. Vesicle numbers represent the experimental chronological order. Although vesicles before aspiration became visibly floppier through time, there is no clear correlation between vesicle number and transition tension.

| $\Delta c$ [mM] | Vesicle number | Tension [mN/m] |
|---|---|---|
| 0 | 1 | 0.0 |
| 0 | 2 | 0.2 |
| 0 | 3 | 0.0 |
| 0 | 4 | 0.0 |
| 0 | 5 | 0.0 |
| 0 | 6 | 0.5 |
| 0 | 7 | 0.0 |
| 0 | 8 | 0.1 |
| 0 | 9 | 0.0 |
| 0 | 10 | 0.0 |
| 0 | 11 | 0.0 |
| 0 | 12 | 0.1 |
| 0 | 13 | 0.0 |
| 0 | 14 | 0.2 |
| 0 | 15 | 0.0 |
| 20 | 1 | 0.9 |
| 20 | 2 | 0.3 |
| 20 | 3 | 0.6 |
| 20 | 4 | 0.4 |
| 20 | 5 | 0.4 |
| 20 | 6 | 0.0 |
| 20 | 7 | 0.0 |
| 20 | 8 | 0.6 |
| 20 | 9 | 1.0 |
| 20 | 10 | 0.1 |
| 20 | 11 | 0.0 |
| 20 | 12 | 0.0 |
| 20 | 13 | 0.0 |
| 20 | 14 | 0.1 |
| 20 | 15 | 0.0 |
| 40 | 1 | 1.9 |
| 40 | 2 | 0.0 |
| 40 | 3 | 1.1 |
| 40 | 4 | 1.4 |
| 40 | 5 | 1.0 |
| 40 | 6 | 0.0 |

**Supplemental Table S3:** Data of vesicle tensions in Figure 5B of the main text. All vesicles had the lipid composition of point 1 in Figure 2A of the main text. Vesicles were electroformed in 100 mM sucrose. To reach concentration differences $\Delta c$ = 20 mM and $\Delta c$ = 40 mM, vesicle solutions were diluted with water. Each vesicle was aspirated by the micropipette, and then the aspiration pressure was slowly increased. Vesicle tension was determined as the tension at which the vesicle first began to deform. This measurement was conducted at room temperature.

| Vesicle number | Time until rupture [s] | Δradius [%] | Ersatz $\Delta c$ [mM] |
|---|---|---|---|
| 1 | 150 | -1.7 ± 2.8 | -5.4 ± 8.9 |
| 2 | 285 | 4.4 ± 3.0 | 12.2 ± 7.4 |
| 3 | 325 | 6.7 ± 3.0 | 17.7 ± 7.0 |
| 4 | 285 | 13.2 ± 3.2 | 31.1 ± 5.8 |
| 5 | 300 | 13.8 ± 3.2 | 32.1 ± 5.8 |
| 6 | 355 | 5.8 ± 3.0 | 15.5 ± 7.2 |
| 7 | 240 | 2.2 ± 2.9 | 6.4 ± 7.9 |
| 8 | 100 | -2.4 ± 2.8 | -7.4 ± 9.1 |
| 9 | 205 | 1.8 ± 2.9 | 5.3 ± 8.0 |
| 10 | 435 | 11.5 ± 3.2 | 27.9 ± 6.1 |
| 11 | 320 | 12.0 ± 3.2 | 28.8 ± 6.0 |
| 12 | 550 | 7.7 ± 3.0 | 20.0 ± 6.8 |
| 13 | 335 | 7.1 ± 3.0 | 18.6 ± 6.9 |

**Supplemental Table S4:** Data corresponding to Figure 5C of the main text. Radii were measured from a time lapse images of a single population of vesicles (Supplemental Movie S1). The lipid composition corresponds to point 1 in Figure 2A of the main text. Additional details are in the caption of Supplemental Movie S1.
